# Multiomics uncovers the epigenomic and transcriptomic response to viral and bacterial stimulation in turbot

**DOI:** 10.1101/2024.02.15.580452

**Authors:** Oscar Aramburu, Belén Gómez-Pardo, Paula Rodríguez-Villamayor, Andrés Blanco-Hortas, Jesús Lamas, Pooran Dewari, Diego Perojil-Morata, Pierre Boudinot, Daniel J. Macqueen, Carmen Bouza, Paulino Martínez

## Abstract

Uncovering the epigenomic regulation of immune responses is essential for a comprehensive understanding of host defence mechanisms, though remains poorly investigated in farmed fish. We report the first annotation of the innate immune regulatory response in the turbot genome (Scophthalmus maximus), integrating RNA-Seq with ATAC-Seq and ChIP-Seq (H3K4me3, H3K27ac and H3K27me3) data from head kidney (in vivo) and primary leukocyte cultures (in vitro) 24 hours post-stimulation with viral (poly I:C) and bacterial (inactive *Vibrio anguillarum*) mimics. Among the 8,797 differentially expressed genes (DEGs), we observed enrichment of transcriptional activation pathways in response to Vibrio and immune pathways - including interferon stimulated genes - for poly I:C. We identified notable differences in chromatin accessibility (20,617 in vitro, 59,892 in vivo) and H3K4me3-bound regions (11,454 in vitro, 10,275 in vivo) between stimulations and controls. Overlap of DEGs with promoters showing differential accessibility or histone mark binding revealed significant coupling of the transcriptome and chromatin state. DEGs with activation marks in their promoters were enriched for similar functions to the global DEG set, but not always, suggesting key regulatory genes being in poised state. Active promoters and putative enhancers were enriched in specific transcription factor binding motifs, many common to viral and bacterial responses. Finally, an in-depth analysis of immune response changes in chromatin state surrounding key DEGs encoding transcription factors was performed. This multi-omics investigation provides an improved understanding of the epigenomic basis for the turbot immune responses and provides novel functional genomic information, leverageable for disease resistance selective breeding.

## INTRODUCTION

The functional annotation of farm animal genomes is important for understanding traits with complex genetic architecture, such as disease resistance, growth, feed efficiency or reproduction (Andersson et al., 2015; Tuggle et al., 2016; Giuffra et al., 2019; Clark et al., 2020). Until recently, functional annotation mainly focused on protein-coding genes using transcriptomics. Transcriptome annotation is now consolidated with robust pipelines (Raghavan et al., 2022) and, throughout the years, transcriptome annotations for human (Frankish et al., 2023), model species (He et al., 2020; Lawson et al., 2020), terrestrial livestock (Summers et al., 2020; Halstead et al., 2021; Overbey et al., 2021) and some aquaculture species (Ramberg et al., 2021; Johnston et al., 2024) have been published.

Non-coding regulatory elements, including promoters, enhancers, silencers and insulators, have been studied in several livestock species but are mostly unexplored in aquaculture species. These elements play essential roles in regulating gene expression and their state can change depending on tissue, cell type, sex, age, and health status (Kellis et al., 2014). Thus, annotation of regulatory elements in different contexts not only aids to address basic questions related to morphology and physiology (Halstead et al., 2020; Pan et al., 2023), but also functional genomic responses to environmental variation (Feinberg, 2007; Ecker et al., 2018; Hu and Barrett, 2017). Genetic variation at non-coding elements also underpins phenotypic variation (Villar et al., 2020; Boltsis et al., 2021), and can thus be leveraged to improve our ability to predict polygenic traits using genomic data (Zhu et al., 2023). In this respect, more than 90% of phenotype-associated single nucleotide polymorphisms (SNPs) identified in human genome-wide association studies (GWAS) are located in non-coding regions (Giral et al., 2018), with similar results reported for livestock (Prowse-Wilkins et al., 2021).

In the past decade, human and livestock functional annotation initiatives have investigated epigenetic mechanisms involved in gene regulation through the study of chromatin state modifications across the genome (Yan et al., 2020a). Chromatin can switch dynamically between active and inactive states in minutes to hours, leaving epigenetic footprints that can be transmitted vertically following DNA replication (Moazed, 2011; Cuvier and Fierz, 2017). Many different sequencing assays have been developed to infer chromatin epigenetic status, including chromatin accessibility (ATAC-Seq; Buenrostro et al., 2015), protein-DNA interactions (ChIP-Seq; Park, 2009) and long-range chromatin interactions (Hi-C; van Berkum et al., 2010). These and other assays are being applied by the FAANG Consortium (Functional Annotation of Animal Genomes; Fang et al., 2019; Foissac et al., 2019; Liu et al., 2020; Clark et al., 2020). Current annotations of chromatin state and regulatory elements remain limited to a few terrestrial farm animal species (Summers et al., 2020, Pan et al., 2021; 2023; Xiang et al., 2021). However, a catalogue of regulatory elements is being generated for several important fish species used in global aquaculture (Johnston et al., 2024), currently the fastest growing animal production sector (Subasinghe et al., 2009; Troell et al., 2023).

Turbot (*Scophthalmus maximus*) is a valuable farmed fish in Europe and Asia (more than 100,000 tons), with the highest production in China (Gao et al., 2023) followed by Spain (APROMAR 2022). Turbot is in its 6^th^ generation of selective breeding and infectious disease outbreaks constitute one of the main challenges this young industry faces (Martínez et al., 2021; Gao et al., 2023). This is a broader trend shared by global aquaculture, where infectious diseases cause losses of totalling more than 5,000 M€ per year (Mishra et al., 2018). Functional annotation of the turbot transcriptome has been performed against high-quality reference genomes (Figueras et al., 2016; Maroso et al., 2018; Xu et al., 2020; Martinez et al., 2021), including for immune-organs stimulated with viruses (Diaz-Rosales et al., 2012), bacteria (Millán et al., 2011; Libran- Pérez et al., 2022) and parasites (Pardo et al., 2012; Robledo et al., 2014; Ronza et al., 2016; Valle et al., 2020). Candidate genes for disease resistance have been further explored by mapping differentially expressed genes (DEGs) within QTL regions (Martínez, 2016; Saura et al., 2019; Aramburu et al., 2023). However, limited attention has been given to non-coding regulatory elements, beyond a recent analysis of chromatin accessibility focussed on early development (Guerrero-Peña et al., 2023). How chromatin state and non-coding regulatory elements are regulated during immune responses remains undefined in turbot and scarcely explored in other farmed finfish.

The head kidney has been targeted in all previous functional genomics studies in turbot investigating pathogen responses (Millán et al., 2011; Díaz-Rosales et al., 2012; Pardo et al., 2012; Robledo et al., 2014; Ronza et al., 2016; Librán-Pérez et al., 2022; Aramburu et al., 2023), due to its central role in fish immunity (Mokhtar et al., 2023). Head kidney is a key lymphoid organ in most marine fishes and, analogous to the mammalian bone marrow, responsible for the production of high leukocyte diversity, including B-lymphocytes, early-stage T-lymphocytes, as well myeloid cells such as granulocytes and monocytes/macrophages (Klosterhoff et al., 2015; Geven et al., 2017; Chen et al., 2022). Innate immunity provides the first line of defence against pathogens and acts following the binding of pathogen-associated molecular patterns (PAMPs) to germline pattern recognition receptors (PRRs), leading to various effector cellular functions targeting pathogen destruction and clearance.

The aim of this study was to generate the first comprehensive functional annotation of the innate immune response of turbot using a chromosome-level reference genome sequence (Martínez et al., 2021). Live fish and primary immune cell cultures were stimulated using mimics of viral and bacterial infections and compared to controls to capture changes in the transcriptome alongside chromatin accessibility and epigenetic state by integrating RNA-Seq, ATAC-Seq and ChIP-Seq data. The experimental design and assays followed the protocols established in the the European Commission Horizon 2020 AQUA-FAANG project (Grant Agreement 817923). We aimed to generate comparable datasets in response to the same bacterial and viral mimics in six commercially important farmed fish species: European seabass (*Dicentrarchus labrax*), gilthead seabream (*Sparus aurata*), rainbow trout (*Oncorhynchus mykiss*), Atlantic salmon (*Salmo salar*), common carp (*Cyprinus carpio*) and turbot (*Scophthalmus maximus*). Our results provide a deeper understanding of the epigenomic basis for innate immunity in turbot and a novel resource to prioritize genetic variation associated with non-coding elements regulating immune responses.

## RESULTS

### Raw sequencing data and sample metadata

A total of 186 multiomic datasets were produced in this study, including RNA-Seq (36), ATAC-Seq (36) and ChIP-Seq (108; 36 per histone mark, plus 6 ChIP-Seq input controls). Full information on samples and metadata is shared in Supplementary tables 1 and 2.

### RNA-Seq

On average, 69,580,246 raw reads per library were produced across the 36 RNA-Seq samples, with 97.2 % mapping to the turbot genome (Supplementary table 3A). Principal Component Analysis (PCA) showed that 77 % of the transcriptome variance was explained by PC1, separating the *in vivo* and *in vitro* stimulations (Figure 1A). Despite all three *in vivo* conditions grouping together, a suggestive spatial segregation was observed mainly across PC1, with the control at one end and the *Vibrio* stimulation at the other. For the *in vitro* stimulations, PC2 clearly separated the *Vibrio* stimulation from poly I:C and control samples, the last two showing some overlap (Figure 1B).

**Figure 1.**
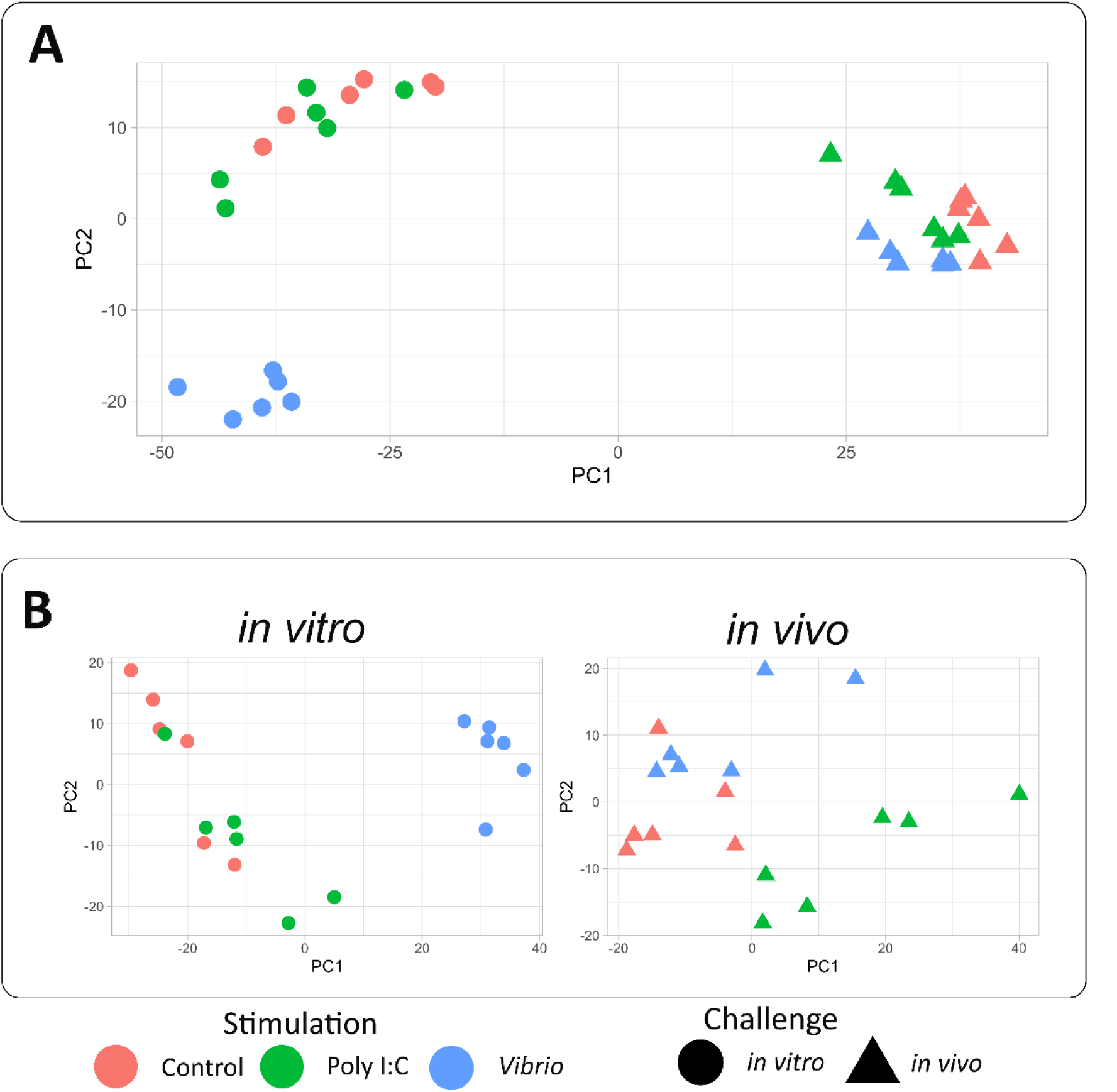
PCA of the *in vivo* and *in vitro* transcriptomic response to stimulation with poly I:C and *Vibrio*. **A)** PCA of the whole transcriptomic dataset; **B)** Independent PCAs for the *in vitro* and *in vivo* transcriptomic assays.

Differential expression analysis was performed by comparing each stimulated condition to the respective controls, both for the *in vitro* and *in vivo* stimulations. In total, 8,797 DEGs were identified across all comparisons (Table 1, Supplementary table 4). For the *in vitro* stimulations, a stronger response was observed for *Vibrio* than poly I:C stimulation, both for up and downregulated genes. Meanwhile, poly I:C showed a higher number of DEGs than *Vibrio* for the *in vivo* stimulations both for up and downregulated genes (Table 1, Supplementary table 4). Overall, more DEGs were detected in the *in vitro* than in the *in vivo* stimulations (7,940 vs 5,758, respectively).

**Table 1.** Number of differentially expressed genes (DEGs) for the *in vitro* and *in vivo* stimulations with *Vibrio* and poly I:C.

| Stimulant | Stimulation | Downregulated DEGs | Upregulated DEGs | Total DEGs |
| --- | --- | --- | --- | --- |
| *Vibrio* | *in vitro* | 3,217 | 3,321 | 6,538 |
|  | *in vivo* | 929 | 910 | 1,839 |
| poly I:C | *in vitro* | 544 | 858 | 1,402 |
|  | *in vivo* | 1,918 | 2,001 | 3,919 |

### Functional enrichment among DEGs

Gene Ontology (GO) analysis identified enriched biological processes for up and downregulated DEGs in all conditions (Supplementary table 5). Among the downregulated DEGs, metabolism, cell cycle and cytoskeleton organization terms were enriched for the *in vitro* stimulations, while a limited number of terms were detected for *in vivo* stimulations (Supplementary table 5).

More abundant and specific enriched GO terms were detected among the upregulated DEGs. Although RNA metabolism was enriched for most stimulations, poly I:C stimulation was linked to a more specific activation of key immune functions (particularly *in vitro*) such as interferon-stimulated genes and cytokine pathways, and regulation of toll-like receptors, besides more general immune terms. DEGs for the *Vibrio in vitro* stimulation were enriched for immune-related terms associated with cytokine and several transport pathways, whereas *Vibrio in vivo* upregulated genes displayed strong enrichment of terms related to cytoskeleton organization, tissue development and syncytium formation (Supplementary table 5).

Comparing upregulated DEGs between conditions, important immune-related GO terms were commonly enriched between both poly I:C and *Vibrio in vitro* stimulations (Figure 2; Supplementary table 6), including interferon type I stimulated genes (*socs1a, socs1b, nod2, nmi*), cytokine signalling (the same genes plus *traf2*, *stat4* and *il15ra*) and major histocompatibility complex type I (MHC-I) pathways (*erap1b*, *tapbpl*, *tapbp*.2). Activation of transcription, protein localization, and a small number of terms associated with the immune-related “peptidyl-arginine modification” (*prmt1, prmt3, prmt5, prmt7*) were commonly overrepresented in poly I:C and *Vibrio in vivo*.

**Figure 2.**
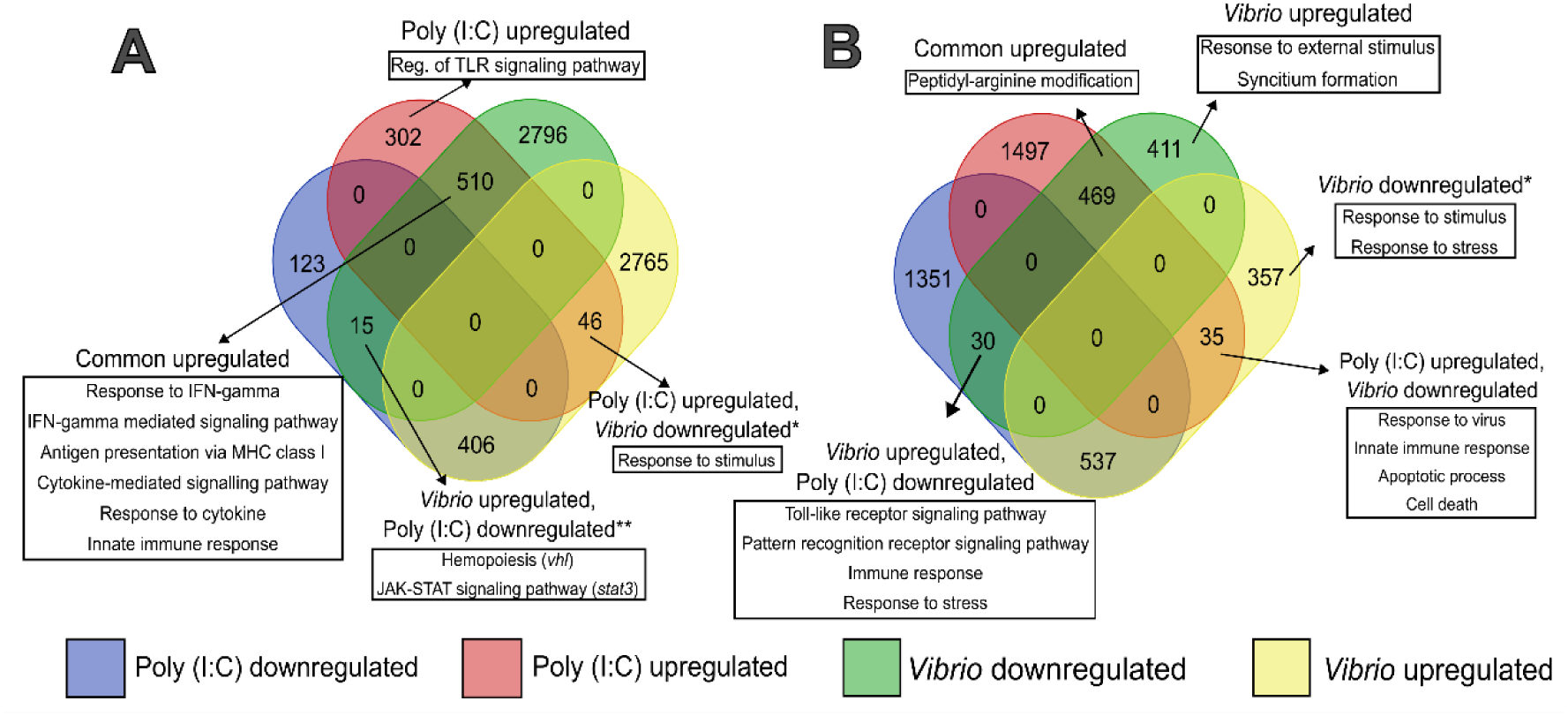
Venn diagrams illustrating overlap of DEGs for: **A)** *in vitro* poly I:C vs *Vibrio*, and **B)** *in vivo* poly I:C vs *Vibrio*. The white boxes show enriched GO terms related to the immune system found in specific subsets. Asterisks (*) highlight cases where suggestive immune-related GO terms were identified, despite not being significantly enriched. Double asterisks (**) highlight specific genes (in parentheses) associated with relevant immune functions. GO terms for the genes in the list were extracted from Ensembl-BioMart.

We detected several DEGs regulated in opposite directions for poly I:C and Vibrio stimulations (Table 2; Figure 2; Supplementary table 6), suggestive of divergent immune response to bacteria and virus. Among the *in vivo* upregulated DEGs for *Vibrio* and downregulated for poly I:C, we observed enrichment of general immune functions, including genes such as *il-1b*, *traf4a* and *f2r*. Conversely, immune-related genes including *nod2*, *sting1*, *irf1b* and *apaf1*, were downregulated for *Vibrio* and upregulated for poly I:C *in vivo*. Interestingly, most genes involved in the type-I IFN response orthologs of human and mouse (interferome database; Rusinova et al., 2023; Clark et al., 2023) were upregulated in the poly I:C stimulations, while downregulated with *Vibrio* extracts, especially in the *in vivo* stimulation (Table 2).

**Table 2.**
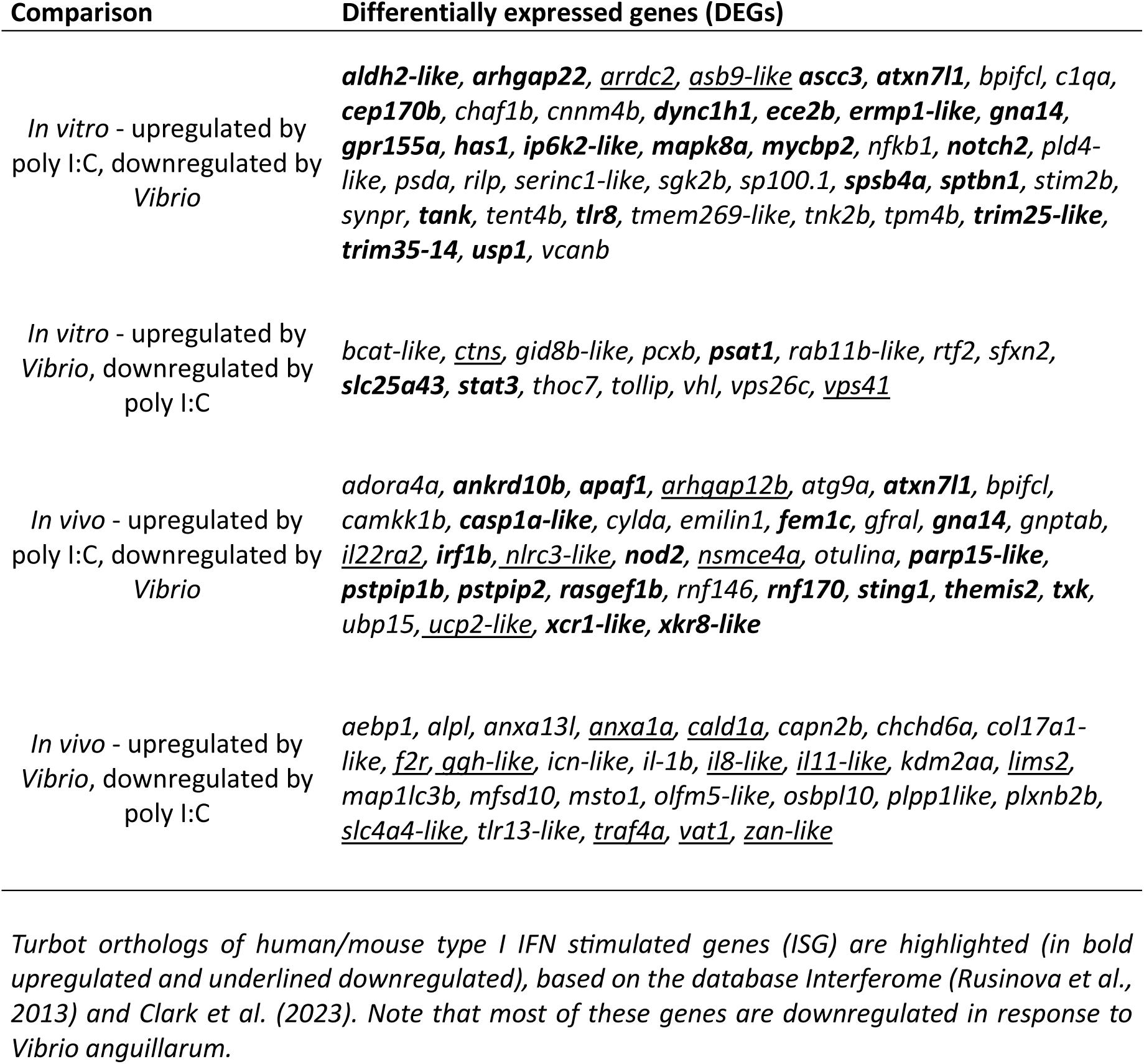
Immune-related DEGs showing opposite responses following *Vibrio* and poly I:C stimulations.

Considering condition-specific DEGs (Figure 2; Supplementary table 6), those upregulated by poly I:C *in vitro* were enriched for terms associated with toll-like receptor signalling pathway (*usp4, tasl, irak3*), while transcriptional activation terms were found among upregulated DEGs for *Vibrio in vitro.* For the *in vivo* stimulations, transcriptional activation GO terms were enriched for upregulated DEGs after poly I:C stimulation, whereas response to stimulus (explained by *tlr3, hamp, tnip1, fxc1a* and *ccr12a*, among other genes) and syncytium formation (explained by *kirrel3l*, *jam2a*, *plekho1b*) were enriched for *Vibrio*.

### ATAC-Seq and ChIP-Seq

Excluding some ATAC-Seq *in vitro* libraries not associated with the type of stimulant, most libraries showed > 95% read mapping to the turbot genome (Supplementary table 3B). On average, we identified 24,251 (*in vitro*) and 62,013 (*in vivo*) peaks per sample for ATAC-Seq; 26,199 (H3K4me3), 13,461 (H3K27ac) and 27,765 (H3K27me3) peaks for the *in vivo* ChIP-Seq data; and 15,870 (H3K4me3), 5,363 (H3K27ac) and 38,784 (H3K27me3) peaks for the *in vitro* ChIP-Seq data. Samples in the PCA plot were mostly segregated across PC1, although PC2 explained most of the variance between histone marks, particularly differentiating repressive (H3K27me3) from active (H3K4me3 and H3K27ac) marks (Figure 3, Supplementary figure 1). *Vibrio* showed the greatest differentiation among the stimulations, while poly I:C and controls were mostly intermingled, particularly for ATAC-Seq (*in vitro* and *in vivo*) and H3K4me3-ChIP-Seq (*in vitro*).

**Figure 3.**
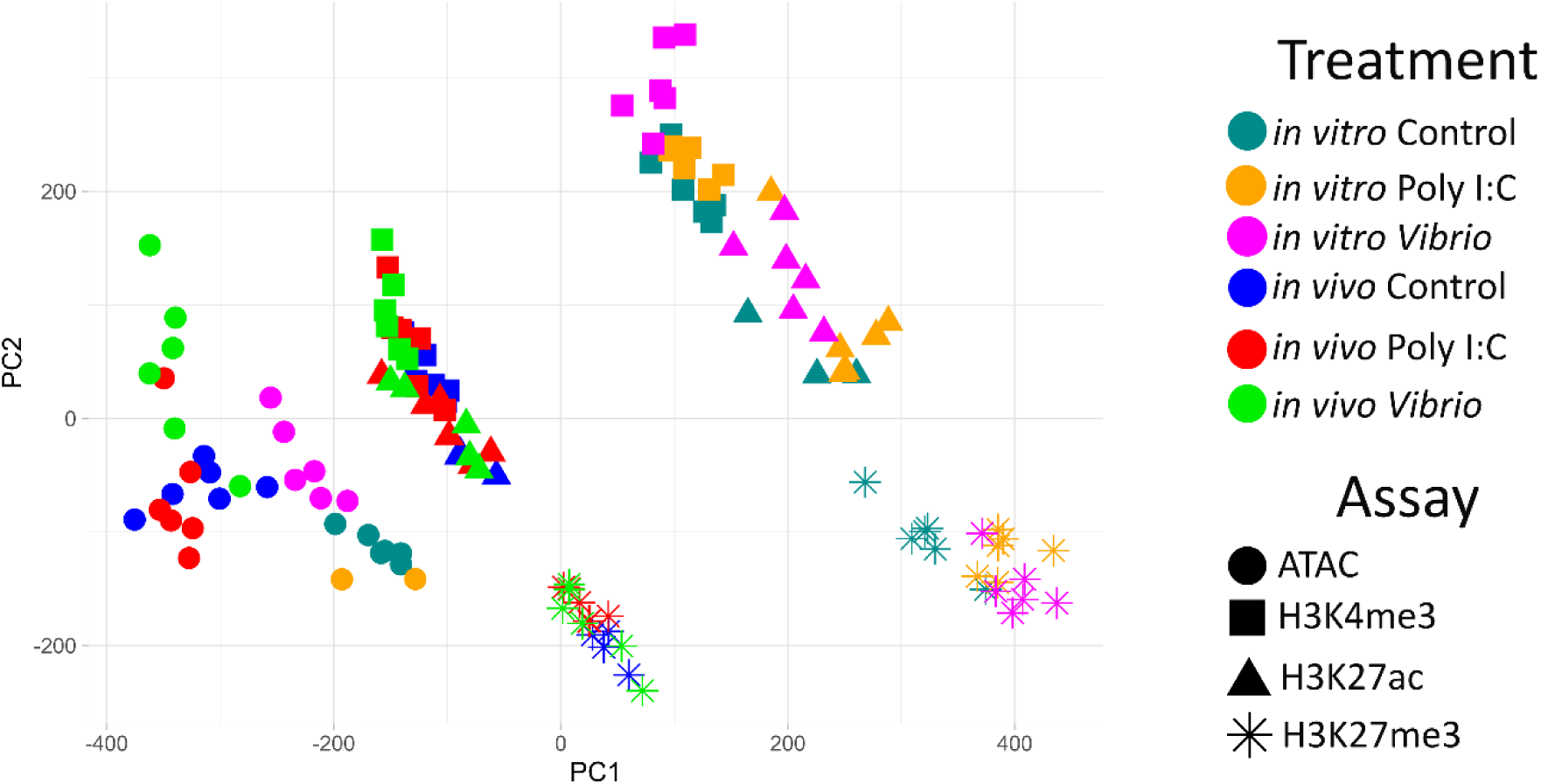
PCA of the *in vivo and in vitro* epigenomic response to stimulation with poly I:C and *Vibrio* for the open chromatin ATAC-Seq and histone ChIP-Seq (H3K4me3, H3K27ac, H3K27me3) samples.

### ChIP-Seq blacklist

Certain genomic regions obscure epigenetic analyses because of anomalous, unstructured, or high signal due to particular genomic features (Amemiya et al., 2019). Using the 21 ChIP-Seq and µChIPmentation input controls (ENA project PRJEB57784) we constructed a blacklist of low confident genomic regions for turbot ChIP assays (Supplementary table 7). On average, 6.98 % (∼39Mb) of the turbot genome was included in the blacklist, consisted of high input signal (5.58%) and low mappability (1.40 %) regions (Supplementary figure 2).

### Chromatin state annotation

Genome-wide chromatin state predictions were produced using ChromHMM employing the ChIP-Seq and ATAC-Seq data from stimulated and control samples from head kidney and leukocytes (Figure 4, Supplementary table 8). For the *in vivo* head kidney samples (Figure 4A), a 10-state model was chosen including promoters/transcription start sites (TSS; States 1, 2, 3 and 4), potential enhancer regions (States 5 and 6), ATAC islands (State 7, i.e. ATAC-peaks lacking histone marks), repressed regions (States 8 and 9) and low signal regions (State 10). For the *in vivo* leukocytes (Figure 4B), an 8-state chromatin model was chosen including promoters/TSS (States 1, 2 and 3), potential enhancer regions (States 4 and 5), ATAC islands (State 6), repressed regions (State 7) and low signal regions (State 8). For each chromatin state map, ±2kb regions around the TSS showed signal specifically for chromatin states 1, 2, 3 and 4 for head kidney (*in vivo*) and states 1, 2 and 3 for leukocytes (*in vitro*), mostly corresponding to promoter regions and/or transcriptionally active regions.

**Figure 4.**
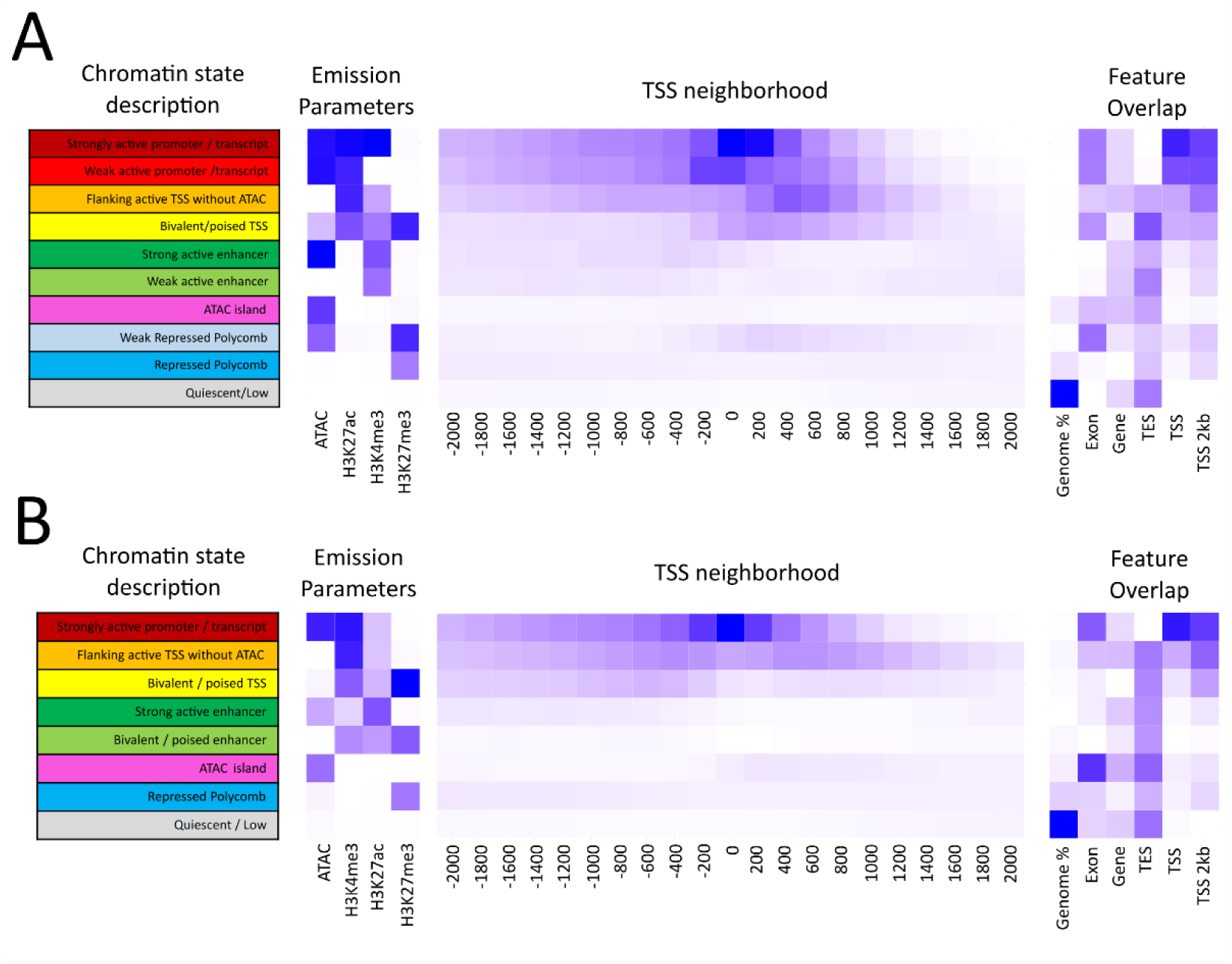
Chromatin state models of the turbot genome: **A)** 10-state model based on *in vivo* samples; **B)** 8-state model based on *in vitro* samples. Heatmaps of the emission parameters for each chromatin state are shown for ChIP-Seq data with three histone marks and ATAC-Seq (left), the neighbourhood around the TSS (middle) and the emission parameters of each chromatin state considered alongside features within the turbot genome (right). TSS: Transcription start site; TES: Transcription end site.

### Differential chromatin accessibility regions (DAR) and differential histone modification regions (DHMR) following stimulation

We next aimed to identify regulatory regions in the turbot genome affected by the immune stimulation. Significant DARs and DHMRs (FDR-adjusted p < 0.05) comparing *Vibrio* and poly I:C stimulations to control samples were identified (Table 3, Supplementary table 9). Globally, a higher number of DARs or DHMRs were detected for up-than for down-regulated regions, more *in vivo* than *in vitro*, and more for *Vibrio* than poly I:C comparisons. Additionally, histone marks H3K27ac and H3K27me3 showed much lower DHMRs than H3K3me3 or ATAC-Seq DARs.

**Table 3.** Differential Accessibility Regions (DARs) and Differential Histone Modification Regions (DHMRs) for the *in vitro* and *in vivo* immune stimulations with viral and bacterial PAMPs.

| PAMP | Stimulation | Assay | Downregulated DAR/DHMR | Upregulated DAR/DHMR | All DAR/DHMR |
| --- | --- | --- | --- | --- | --- |
| *Vibrio* | *in vitro* | ATAC | 2,739 | 17,878 | 20,617 |
|  |  | H3K4me3 | 699 | 10,755 | 11,454 |
|  |  | H3K27ac | 2 | 623 | 625 |
|  |  | H3K27me3 | 6 | 1,211 | 1,217 |
|  | *in vivo* | ATAC | 43,159 | 16,733 | 59,892 |
|  |  | H3K4me3 | 287 | 9,988 | 10,275 |
|  |  | H3K27ac | 0 | 3 | 3 |
|  |  | H3K27me3 | 4 | 0 | 4 |
| poly I:C | *in vitro* | ATAC | 0 | 0 | 0 |
|  |  | H3K4me3 | 0 | 0 | 0 |
|  |  | H3K27ac | 0 | 20 | 20 |
|  |  | H3K27me3 | 0 | 0 | 0 |
|  | *in vivo* | ATAC | 25 | 15 | 40 |
|  |  | H3K4me3 | 0 | 1,036 | 1,036 |
|  |  | H3K27ac | 0 | 38 | 38 |
|  |  | H3K27me3 | 126 | 0 | 126 |

### Association of ATAC-Seq and ChIP-Seq data with RNA-Seq data

We tested if DARs and DHMRs for H3K4me3 and H3K27ac marks annotated as promoter/TSS regions (up to -1kb upstream of TSSs) corresponded to DEGs under the same experimental conditions (Hypergeometric distribution test, p < 0.05). DARs and DHMRs were much more overrepresented at the promoter regions of up-rather than down-regulated DEGs (Table 4, Supplementary table 10), suggesting changes in chromatin state associated with the activation of genes. We performed GO enrichment analyses of those upregulated DEGs within each experimental condition t (P < 0.05, Table 4). Significant enrichment (FDR < 0.05) included several metabolic activities and particular immune functions, including antigen processing/presentation and apoptotic/cell death pathways (Supplementary table 11). Specifically, enriched terms were mostly associated with RNA processing (in particular tRNA, rRNA, ncRNA), ribosome biogenesis, and translation for the *Vibrio in vitro* stimulation, whereas for protein localization to nucleus and organelle, DNA replication and carbohydrate metabolism for the *Vibrio in vivo* stimulation. The term ‘peptidyl-arginine modification’ was also enriched involving *prmt1, prmt3, prmt5* and *prmt7* genes, as outlined before for the whole DEG analysis (RNA-Seq). The poly I:C *in vivo* stimulation showed enrichment in immune-related processes, including ‘antigen processing and presentation via MHC-I’ (explained by *erap1b, tapbpl*) and ‘regulation of programmed cell death’ (explained by *apaf1, bida, casp8 and 10, socs3a, grinab, bcl2l10*; Supplementary table 11).

**Table 4.**
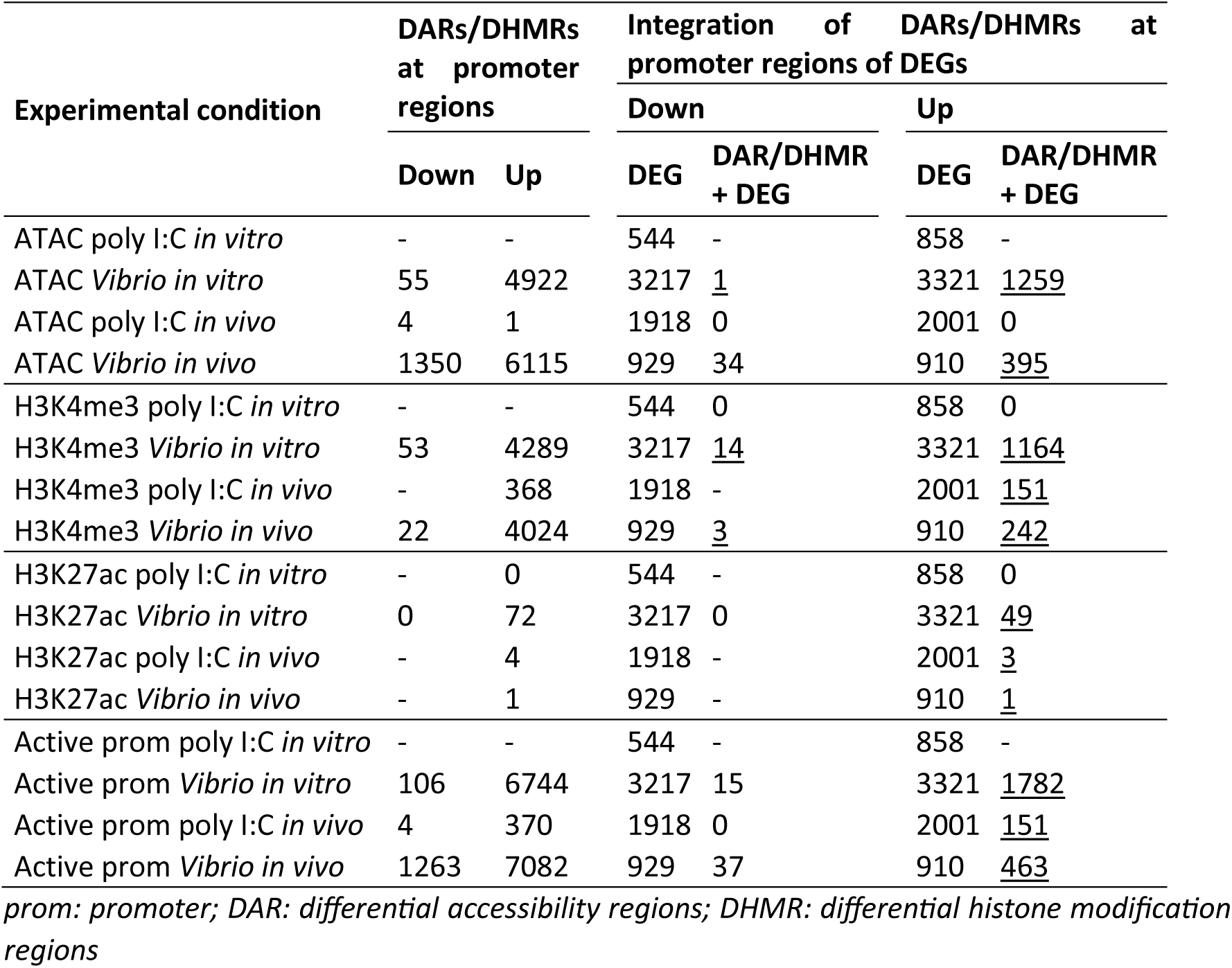
Overlap of promoter DARs and DHMRs with DEG promoters (Hypergeometric test, p < 0.05). Significant results are underlined. Refer to Supplementary table 10 for further details.

| Experimental condition | DARs/DHMRs at promoter regions |  | Integration of DARs/DHMRs at promoter regions of DEGs |  |  |  |
| --- | --- | --- | --- | --- | --- | --- |
|  |  |  | Down |  | Up |  |
|  | Down | Up | DEG | DAR/DHMR + DEG | DEG | DAR/DHMR + DEG |
| ATAC poly I:C *in vitro* | - | - | 544 | - | 858 | - |
| ATAC *Vibrio in vitro* | 55 | 4922 | 3217 | <u>1</u> | 3321 | <u>1259</u> |
| ATAC poly I:C *in vivo* | 4 | 1 | 1918 | 0 | 2001 | 0 |
| ATAC *Vibrio in vivo* | 1350 | 6115 | 929 | 34 | 910 | <u>395</u> |
| H3K4me3 poly I:C *in vitro* | - | - | 544 | 0 | 858 | 0 |
| H3K4me3 *Vibrio in vitro* | 53 | 4289 | 3217 | <u>14</u> | 3321 | <u>1164</u> |
| H3K4me3 poly I:C *in vivo* | - | 368 | 1918 | - | 2001 | <u>151</u> |
| H3K4me3 *Vibrio in vivo* | 22 | 4024 | 929 | <u>3</u> | 910 | <u>242</u> |
| H3K27ac poly I:C *in vitro* | - | 0 | 544 | - | 858 | 0 |
| H3K27ac *Vibrio in vitro* | 0 | 72 | 3217 | 0 | 3321 | <u>49</u> |
| H3K27ac poly I:C *in vivo* | - | 4 | 1918 | - | 2001 | <u>3</u> |
| H3K27ac *Vibrio in vivo* | - | 1 | 929 | - | 910 | <u>1</u> |
| Active prom poly I:C *in vitro* | - | - | 544 | - | 858 | - |
| Active prom *Vibrio in vitro* | 106 | 6744 | 3217 | 15 | 3321 | <u>1782</u> |
| Active prom poly I:C *in vivo* | 4 | 370 | 1918 | 0 | 2001 | <u>151</u> |
| Active prom *Vibrio in vivo* | 1263 | 7082 | 929 | 37 | 910 | <u>463</u> |

*prom*: promoter; *DAR*: differential accessibility regions; *DHMR*: differential histone modification regions

### Transcription factor binding motif (TFBM) analysis

To establish TFs potentially associated with chromatin state regulation following immune stimulation, the enrichment of TFBMs within DAR/DHMRs annotated as promoter or enhancer regions were examined. Significant enrichment was detected for poly I:C *in vivo* and for *Vibrio in vitro* and *in* vivo stimulations. On average across stimulations, 4 (range: 0-11) and 71 (range: 0-132) enriched TFBMs were detected for down- and upregulated promoters, respectively, while 14 (range: 6-27) and 49 (range: 1-80) respective TFBMs were enriched for downregulated and upregulated enhancers (Supplementary table 12).

Most TFBMs and associated TFs predicted for active promoters and enhancers were enriched in several stimulations: 68 TFs for both *Vibrio* stimulations and *in vivo* poly I:C; 59 TFs for both *Vibrio* stimulations; and 20 TF for both *in vivo* stimulations (Figure 5; Supplementary table 12).

**Figure 5.**
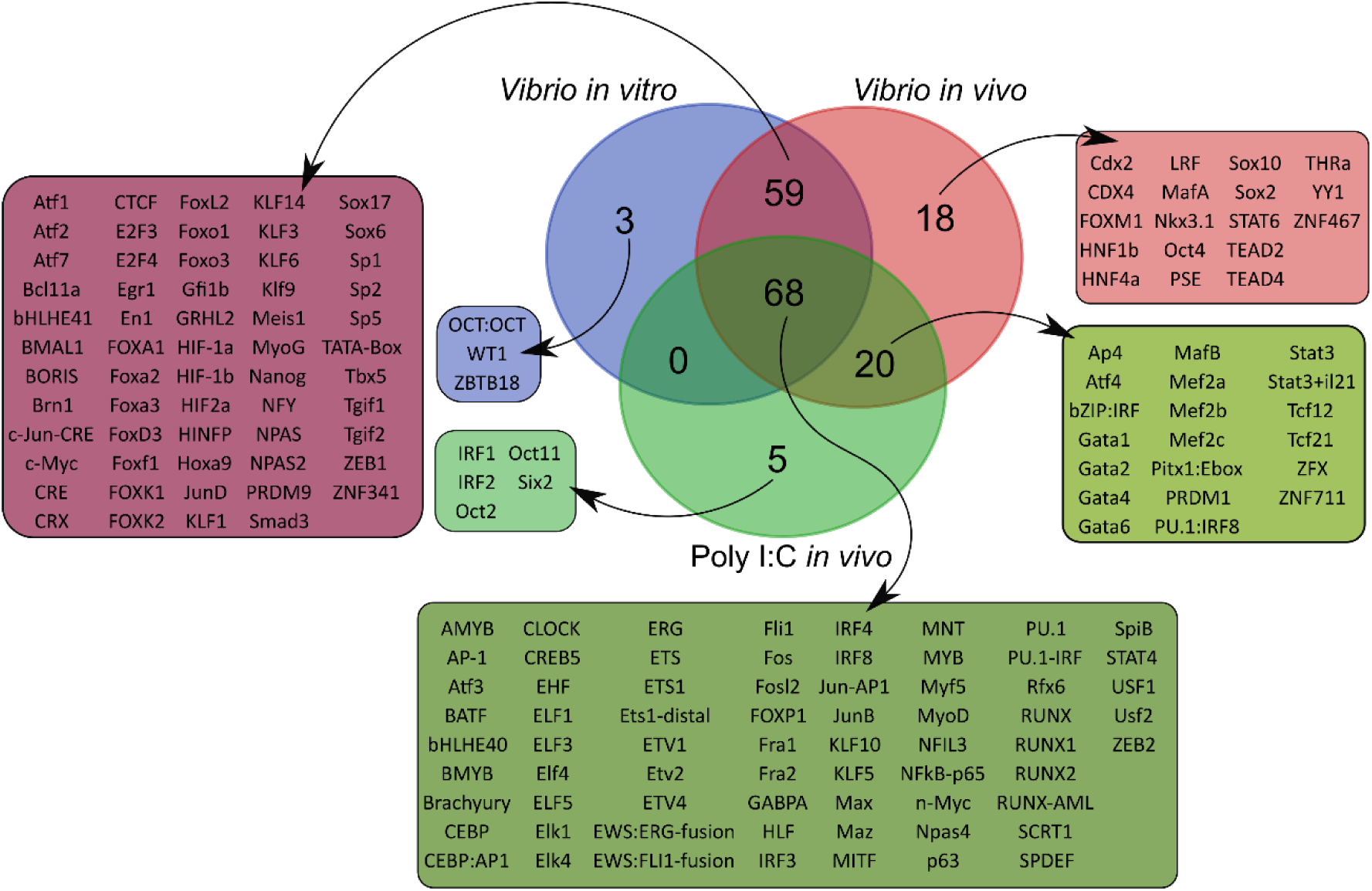
Venn diagram illustrating overlap of enriched transcription factor binding motifs (TFBMs) in upregulated promoter and enhancer regions in response to the different stimulants. Each coloured box shows the TFBM found.

To further investigate the functions of TFs with enriched motifs, we used the Molecular Complex Detection algorithm (MCODE) and protein-protein interaction data (PPI) to identify connected functions among differentially expressed TFs and among TFs with DARs or DHMRs on their promoters (Supplementary figures 3 and 4, Supplementary table 13).

For the *Vibrio in vitro* stimulation, interconnected clusters of TFs associated with hemopoiesis and immune functions, particularly myeloid leukocyte differentiation, FGF signalling pathway, regulation of neutrophil differentiation, lymphocyte activation, apoptosis and the ERK1/2 immune signalling, were identified. For the *Vibrio in vivo* stimulation, we detected clusters of TFs associated with similar functions (e.g. hemopoiesis, the ERK1/2 signalling pathway, myeloid cell development and cell death), but also other functions such as insulin signalling, regeneration, and oestrogen-dependent expression (Supplementary figures 3 and 4). Finally for the poly I:C *in vivo* stimulation, myeloid leukocyte differentiation, regulation of cell differentiation, cell morphogenesis and biosynthesis and growth showed connected TF subsets.

Using the Integrative Genomics Viewer (IGV), we visualized the variation in chromatin states around the promoters of 12 TF-DEGs selected from Supplementary table 13 to show the variance in chromatin state distribution across TFs and their regulatory elements following activation by the different stimulants (Figure 6; Supplementary Figure 5).

**Figure 6.**
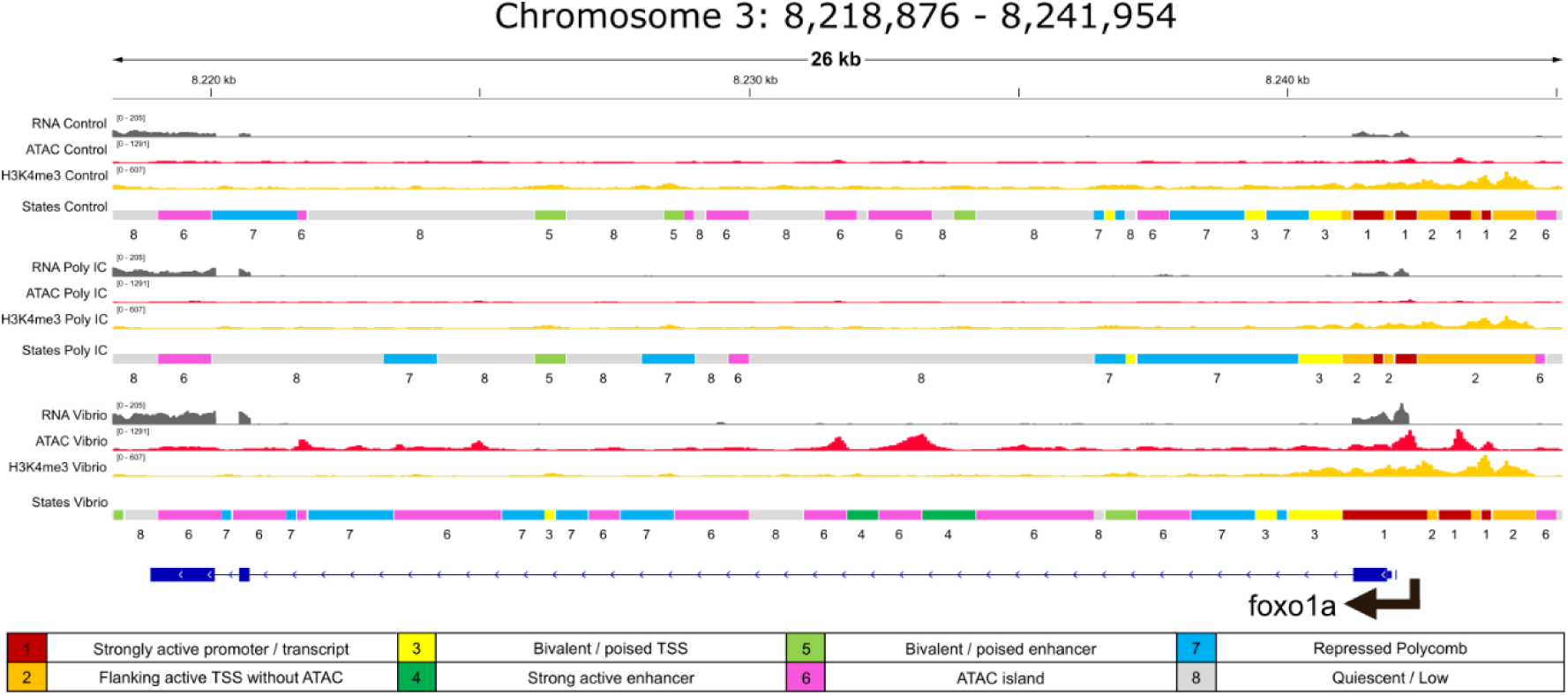
Chromatin state predictions and RNA/ATAC/ChIP-Seq tracks in the transcription factor coding gene *foxo1a*, which was upregulated and showed DAR around the promoter for *Vibrio in vitro* response. Transcription factor binding motifs for FOXO1A were also enriched in the promoter DARs of DEGs upregulated in response to *Vibrio in vitro*.

## DISCUSSION

This study represents the first epigenomic analysis of the turbot head kidney, the primary hematopoietic and lymphoid organ of teleosts, in response to viral (poly I:C) and bacterial (inactivated *V. anguillarum*) mimics. While head kidney transcriptomic responses to bacteria, virus and parasites have been extensively investigated in turbot (Pardo et al., 2008; Millán et al., 2010; Domínguez et al., 2013; Díaz-Rosales et al., 2012; Robledo et al., 2014; Ronza et al., 2020; Valle et al., 2020; Libran-Pérez et al., 2022), the epigenetic regulation of chromatin states following immune stimulation has not been explored before. The head kidney response was explored through intraperitoneal injection, reflecting the response in the whole-body, including interactions among immune organs, and through *in vitro* stimulation of primary leukocyte cultures, reflecting their direct interaction with stimulants. The use of the same viral and bacterial mimics will enable further comparative evaluation among the six fish species included in the AQUA-FAANG project pertaining to five teleost orders (Salmoniformes, Cyprinidontiformes, Spariformes, Perciformes and Pleuronectiformes). We first discuss the transcriptional response to immune stimulations, which provide context to the chromatin assays performed to understand how transcription is regulated from transcription factors up to immune genes in the turbot.

### Transcriptomic response to immune stimulation

Upregulated genes with key immune roles, such as interferon and cytokine pathways and toll-like receptor regulation, were found in both poly I:C stimulations and in *Vibrio in vitro* stimulation. Interferons (IFNs) are a subset of class II cytokines with crucial roles in immune response, especially against viruses but also intracellular bacteria (Gan et al., 2019). Interestingly, *tlr3*, encoding the toll-like receptor 3, which induces IFN production after interaction with poly I:C (Kumar et al., 2006), was upregulated in the *Vibrio in vivo* stimulation but not differentially expressed in poly I:C *in vivo* and downregulated after both *in vitro* stimulations. Signal exhaustion offers a plausible explanation for *tlr3* downregulation *in vitro*, with the 24-hour post-injection (hpi) sampling time missing the immediate interaction between leukocytes and stimulants. However, we cannot discard alternative hypotheses related to changes or diversification of the TLR3 signalling pathway in flatfish or specifically in turbot, as suggested here by *tlr3* activation in response to *Vibrio in vivo* stimulation and elsewhere to the parasite *P*. *dicentrarchi* (Figueras et al., 2016). In fact, we identified other interferon-stimulated DEGs in poly I:C both *in vitro* and *in vivo*, including *socs1a* and b (suppressor of cytokine signalling 1), *nod2* (nucleotide binding oligomerization domain containing 2) and *nmi* (N-Myc interactor), typically participating in type I IFN responses (Blumer et al., 2017; Ahn et al., 2021). *socs1* is an inducible negative regulator of IFN (Blumer et al., 2017), a mechanism to maintain tissue homeostasis following the IFN response; *nod2* encodes for an intracellular receptor triggering innate antibacterial and antiviral signalling pathways, including IFN (Nie et al., 2017); and *nmi* increases STAT-mediated transcription in response to IFN-gamma (Zhu et al., 1999). The activation of genes involved in positive and negative regulation of IFN-signalling suggests a fine adjustment to an intense IFN response to recover tissue homeostasis (Ivashkiv and Donlin, 2014; Murira and Lamarre, 2016). Also, the upregulation of these four key genes after *Vibrio in vitro* stimulation is consistent with past reports that place IFN regulation as a key point of the immune response (Kopitar-Jerala, 2017). In this regard, comparative analysis in the AQUAFAANG project will enlighten conserved and specific mechanisms of immune response regulation across different farmed fish lineages. Future studies in turbot should explore shorter periods following *in vitro* stimulation to provide further understanding of the head kidney response to viral and bacterial mimics, as done in other species (Hasanuzzaman et al., 2020; Herrera-Uribe et al., 2020).

Upregulated DEGs shared by both *in vivo* stimulations were involved in the activation of transcription and translation and protein modification/localization. This included four members of the immune-related protein arginine methyltransferase family (*prmt1*, *prmt3*, *prmt5* and *prmt7*), enriched under the term “peptidyl-arginine modification”. *prmt* family members regulate transcription and translation and are also involved in signal transduction during inflammation and response against poly I:C and bacterial lipopolysaccharide (LPS; Srour, 2022). These four genes were also activated in both *Vibrio* stimulations, suggesting an important role in the response to *Vibrio*.

We also verified if immune genes could be regulated in opposite directions after stimulation with viral and bacterial mimics, which could reflect antagonistic immune responses. Increased susceptibility to bacterial superinfections induced by innate antiviral responses has been reported in several models in mammals (Barman and Metzger, 2021) in type I and type II IFN responses (Navarini et al., 2006; Sun and Metzger, 2008). Such mechanisms are likely present in teleosts due to the high conservation of these pathways, and in fact, different patterns of resistance to viral and bacterial infections were observed among isogenic rainbow trout lines (Fraslin et al., 2020). However, no negative correlations for resistance to bacteria and viruses has been observed in the few studies addressing this issue in relation to selective breeding (Ødegård et al., 2007; 2011; Bangera et al., 2011). Indeed, understanding the cellular basis of antagonistic responses to viruses and bacteria in fish will require more research, but here we identified examples of opposite gene expression responses, including genes from the core type I IFN response, conserved between teleosts and humans (Levraud et al., 2019; Clark et al., 2023), such as *sting1* (stimulator of interferon response CGAMP interactor 1, Zhang et al., 2021*), irf1b* (interferon regulatory factor 1; Sullivan et al., 2021) and *nod2* (mentioned above), downregulated by *Vibrio* and upregulated by poly I:C *in vivo*. Reciprocally, the critical pro-inflammatory cytokine *il1b* (interleukin 1 beta, Sahoo et al., 2011) and the regulator of inflammation *traf4a* (TNF receptor associated factor 4, Marinis et al., 2012) were upregulated by *Vibrio* but downregulated by poly I: C. Interestingly, a majority of genes induced by poly I:C and repressed by *Vibrio* extracts had human / mouse ISG orthologs (22/42 *in vitro*, 18/34 *in vivo*); in contrast, many genes induced by *Vibrio* extracts and repressed by poly I:C, especially *in vivo,* had human/mouse orthologs downregulated by type I IFN. Overall, these observations indicate that a significant part of these contrasted responses is mediated by genes functionally conserved between fish and mammals.

### Epigenetic assays and their association with transcriptomic response

Activation and recruitment of transcription factor (TF) subsets is at the top of the specific transcriptome cascade response (Alvarez et al., 2020; Weidemüller et al., 2021). In our study, the exploration of promoters of differentially expressed TFs, also involving differential accessibility regions (DARs) and differential histone modification regions (DHMRs), barely showed changes in the chromatin state distribution between treatments (Supplementary Figure 5). This is the case for instance of egr*1* (early growth response 1), which plays a key role in cell survival, proliferation and cell death, and regulates the expression of *il1b* and *cxcl2* (CXC motif chemokine ligand 2). The same was observed for *meis1* (myeloid ecotropic viral integration site homeobox 1), related to bone marrow hematopoiesis in mammals, and *zeb2* (zinc finger e-box binding homeobox 2), associated with macrophage infiltration (Taylor et al., 2022). In all these examples, the TF genes showed bivalent/poised promoters in the three *in vivo* samples (control, *Vibrio* and poly I:C). A similar situation was found *in vitro*, where promoters of genes, such as *irf3* (interferon regulatory factor 3) and *mitf* (melanocyte inducing transcription factor), showed activation signals regardless of the experimental condition. In previous studies, these bivalent/poised states have been interpreted as a mechanism allowing rapid responses, e.g. for genes induced during the pro-inflammatory response (Herrera-Uribe et al., 2020; Martínez de Paz and Josefowicz, 2021) or during mammal embryonic development and in germ cells (Lesch and Page, 2014; Barbieri et al., 2017). Thus, the expression of primary response genes, such as the TF genes exemplified above, may be allowed by permissive chromatin states.

The presence of differential activation signals at promoter regions, either in the form of open chromatin regions or the histone marks H3K4me3 and H3K27ac, are associated with differential gene expression (Stępniak et al., 2021). H3K27ac is widely accepted as one of the most dynamic activation marks in eukaryotes (Saeed et al., 2014). Interestingly, we did not find important changes between experimental conditions in our study, where H3K4me3 was the most dynamic among the ChIP-Seq marks. We suspect that the increase in H3K4me3 could also be an effect of sampling being conducted 24 hours post inoculation (hpi)pi, especially for the *in vivo* stimulations. Previous studies have reported that H3K4me3 may not have an active role in activating transcription, an effect that should be detectable early after stimulation, but instead in marking transcriptional activity itself, suggesting that H3K4me3 modification may have a role in maintaining transcriptional consistency or memory of previous states (Howe et al., 2017).

DEG promoters overlapped with DAR/DHMRs for upregulated genes in most conditions (Table 4). GO enrichment of DEGs with activation signals at promoters (Supplementary table 11) was similar to that observed with the whole DEG dataset (Supplementary table 5). *Vibrio* stimulations were particularly enriched in terms associated with transcription activation, but also with some immune functions, while poly I:C (especially *in vivo*) showed a more specific response, enriched in immune-related processes (Supplementary table 6). However, DEGs associated with some key immune functions in the transcriptomic analysis, such as response to cytokine stimulus, did not show DAR/DHMRs, suggesting that these gene promoters could be already accessible before stimulation. Cytokines are one of the core initiators of the inflammatory response (Secombes et al., 2001), thus a bivalent state, ready for activation or inactivation of expression, might explain this observation (Figure 5).

Regardless of predicted chromatin states around gene promoters, we observed an increased expression of immune-related genes following poly I:C and Vibrio stimulation suggesting further chromatin unpacking (Table 3), even if chromatin state is not strongly affected. In fact, many differential regions were detected between stimulations, particularly DARs (i.e. ATAC-Seq) and DHMRs (for H3K4me3) for *Vibrio*. In comparison, fewer DARs and DHMRs were detected for the poly I:C stimulations. We interpret these observations as a return, both in vivo and in vitro, to the native chromatin state after 24 hpi, which may suggest the initial response at this point is exhausted, as suggested by the transcriptomic data.

Predicted TFBMs enriched within DARs/DHMRs in the promoters and enhancer-state regions predicted by ChromHMM of upregulated DEGs, provided a more detailed picture of regulatory elements changed by the immune stimulations (Figure 5, Supplementary table 12). Most of the enriched TFBMs were shared between at least two treatments, which supports their general regulatory function. Among those shared between most stimulations (excluding poly I:C *in vitro*), we found several TFBMs of the ETS-family, which are involved in cell proliferation, apoptosis and lymphocyte development (Watson et al., 2002). We also found TFBMs enrichment of PU.1, a master TF for the myeloid lineage, that promotes chromatin accessibility, also identified in a similar study in pig (Turkistany and DeKoter, 2011; Chen et al., 2020; Herrera-Uribe et al., 2020). PU.1 is thought to promote binding of other TFs that were enriched in our study, including AP1. This protein is activated by TLRs during the immune response (Shan et al., 2018) and regulates gene expression in response to cytokines, stress, and bacterial and viral infections (Kim et al., 2014), and notably the IFN regulatory factors (IRFs, particularly IRF3, IRF4 and IRF8). The IRFs are key regulators of innate antiviral and antibacterial responses in vertebrates including fish (Clark et al., 2021; Han et al., 2023). The observed ontology enrichment clustering of these TFs reinforces the similarities between the conditions in our study (Supplementary figures 3 and 4).

To illustrate the functionality of our turbot epigenomic atlas, we explored the chromatin state changes in a selection of 12 TF genes (Figure 6, Supplementary Figure 5). These genes were differentially expressed and showed DARs/DHMRs in their promoters. Immune-responsive chromatin state remodelling was found, for instance, for *irf8* (interferon regulatory factor 8) in the *Vibrio in vitro* stimulation. Here, an extension of the “strongly active promoter” state around the TSS region was visible compared to the non-stimulated and poly I:C stimulated samples, alongside an overall expansion of the “strongly active transcript” and “ATAC island” states downstream. IRF8 is a key regulator of the NF-κB signalling pathway during inflammation, along with IRF3 (Yan et al., 2020b). Both *irf* genes were differentially expressed in poly I:C and *Vibrio in vitro* stimulations (*irf8* also in poly I:C *in vivo*) and showed promoter DARs in *Vibrio* stimulations. Interestingly, chromatin state changes were barely observed in the vicinity of the *irf3* promoter (Supplementary Figure 5).

Another example was found in *Vibrio* and poly I:C *in vivo* stimulations for *bcl11a* (BCL11 transcription factor A), a key regulator of dendritic cell differentiation (Ippolito et al., 2014) and negative regulator of p53 (Yu et al., 2012). This gene was differentially expressed in both stimulations, but only showed a DAR promoter in response to *Vibrio* stimulation. The promoter/TSS region was annotated by ChromHMM as “weak repressed Polycomb” in the control, which changed in both the *Vibrio* and poly I:C stimulations, extending to “bivalent/poised” state, and in the case of *Vibrio* showing an extension to “Strongly active promoter/transcript” state. Weak and strong signals of active enhancers were also detected in the poly I:C and *Vibrio* stimulations within the second intron of the *bcl11a*, that were not present in the controls (Supplementary Figure 5). A similar situation was found *in vitro* for *foxo1a* (Forkhead box O1), a TF-coding gene that participates in mucosal (innate) immune response regulating the expression of antimicrobial peptides and promoting phagocytosis during bacterial and parasitic infections (Cabrera-Ortega et al., 2017; Graves and Milovanova, 2019). This gene was differentially expressed in *Vibrio in vitro* and showed an extension of the “Strongly active promoter” state compared to the control and specially the poly I:C stimulation, while conserving the “bivalent/poised TSS” stretches downstream of the promoter. This was followed by many “ATAC islands” along the gene that were also present, although more scarcely, in the control, and almost absent in poly I:C. Finally, a “medium enhancer” state was detected within the first intron, present only in the *Vibrio* condition. These observations highlight the relevance of epigenetic chromatin marks at the first intron and other intronic regions associated with gene expression modulation (Jo and Choi 2019; Johnston et al. 2019), as target regulatory annotations to explore functional variants for disease resistance and selective breeding.

### Conclusions and perspectives

In summary, our study provides the first atlas of regulatory elements in turbot head kidney and leukocytes during the early response to viral and bacterial stimulation, as a contribution to the AQUA-FAANG project within the umbrella of the FAANG initiative. The integration of ATAC-Seq and ChIP-Seq data suggests that changes in chromatin state distribution were not as frequent between stimulations and controls as expected. However, the presence of DARs and DHMRs between the stimulations and the controls, which broadly overlapped with DEGs, provides clear evidence for gene regulation at the epigenetic level that underpins changes in gene expression driving immune functions. Overall, this epigenomic atlas will help to decode the molecular mechanisms underlying turbot immune responses to viral and bacterial stimuli and offers a novel resource for developing selective breeding strategies for controlling diseases, one of the main concerns of turbot industry. Future work will benefit from linking the regulatory annotations generated in this study with genetic variants defined by whole genome re-sequencing (Johnston et al., 2024), to help prioritize causal genetic variants for disease resistance traits underpinned by gene expression responses to pathogens.

## METHODS

### Animals

Thirty 8-month-old immature turbot specimens provided by Stolt Sea Farm SA (Ribeira, Spain) were housed in indoor tanks with recirculating seawater at the facilities of the Aquarium of the University of Santiago de Compostela (Spain) for a period of acclimation of 15 d at 16 °C (Supplementary table 1). All fish were fasted for 24 h before stimulations were performed. Eighteen fish were stimulated *in vivo* by intraperitoneal injection, while the other 12 were used for leukocyte isolation for *in vitro* stimulation (see following sections). Fish were anesthetized by bath (MS-222; 100mg/L) and then euthanized by anaesthetic overdose (MS-222; 150 mg/L) before tissue sampling. All animal procedures were approved by the Bioethics Committee of the University of Santiago de Compostela (body authorized according to R.D. 53/2013) and with the authorization of the Xunta de Galicia Regional Government.

### Protocols

Detailed protocols for the *in vivo* and *in vitro* stimulations, RNA isolation, ATAC-Seq and ChIP-Seq (including library preparation) followed for turbot are available in the FAANG repository (data.faang.org; URLs for protocols in Supplementary tables 1 and 2).

### *In vivo* immunostimulation

Six fish were used per experimental condition for the *in vivo* stimulations: i) poly I:C for viral mimic immunostimulation; ii) killed *Vibrio anguillarum* for bacterial immunostimulation; and iii) phosphate buffered saline (PBS) for control. For poly I:C (Sigma P1530), we prepared a working stock at 5 mg/ml in PBS, preheated to 55 °C (15 min) and cooled at room temperature (20 min) before use. Fish were then injected intraperitoneally with 5 µg per g fish weight. For bacterial immunostimulation, an extract of *V. anguillarum* (strain P0382; INRA, France) was used. Bacteria were cultured in a tryptic soy broth medium to an OD600 (optical density at 600 nm) of 1.5. The bacterial pellet (derived from 100 ml of full-grown culture) was washed in an isotonic solution of NaCl (9 g/L) four times and resuspended in 1 ml of the same solution. Bacteria were killed by incubation for 30 sec at 100 °C, allowed to cool at room temperature and stored at -80 °C. The bacteria extract was inoculated in each specimen, diluted in PBS (1:10) for a final volume of 100 µl. Control fish were injected with 100 µl of PBS. After 24 h, head kidney samples were extracted, washed with PBS and cut into at least three pieces (> 20 mg each); two were flash frozen on dry ice for ATAC-Seq and ChIP-Seq and the other was immersed in RNAlater (Thermofisher Scientific) for RNA extraction and RNA-Seq. All samples were then stored at -80°C (Figure 7A; Supplementary table 2).

**Figure 7.**
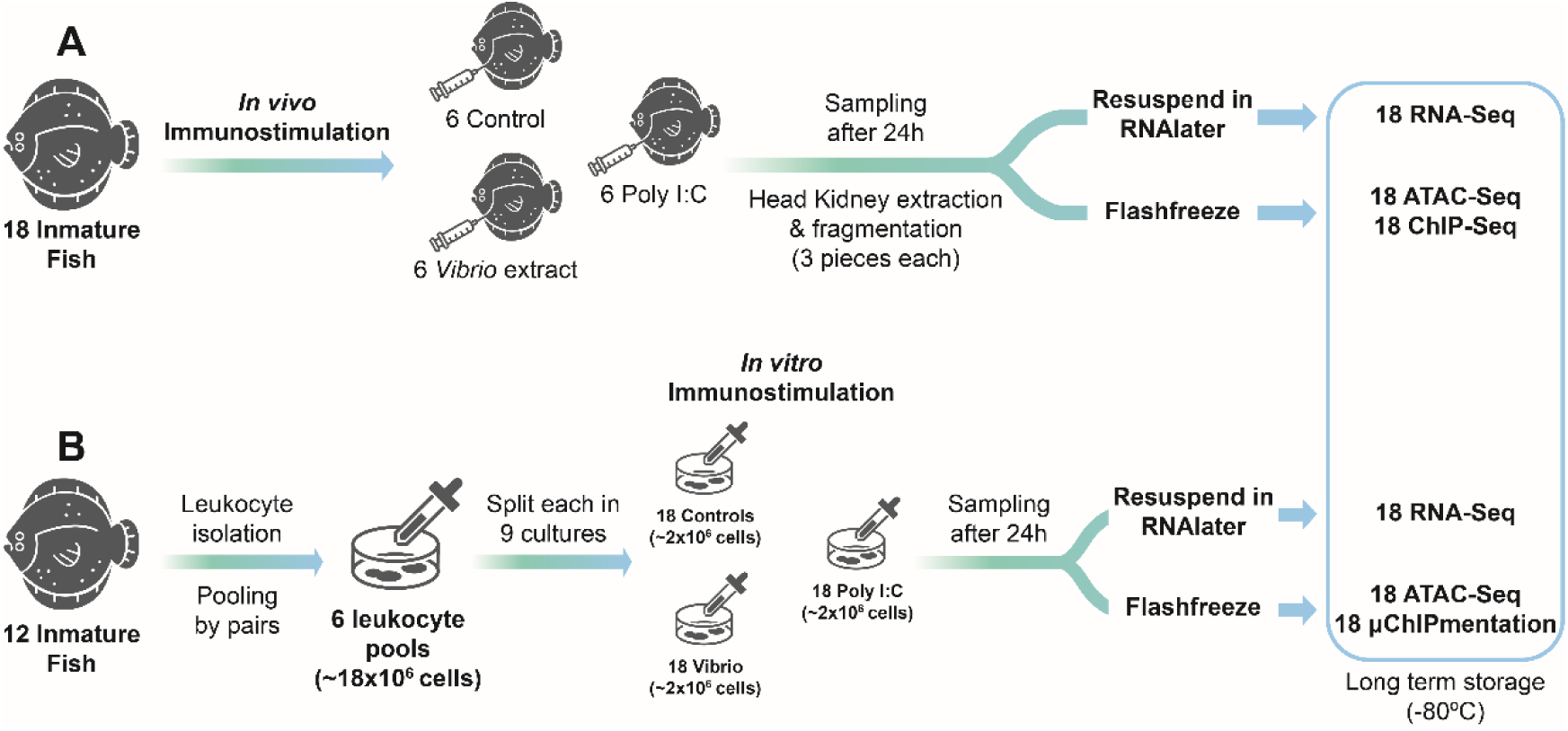
Experimental design followed for turbot immunostimulations: **A)** *in vivo* immunostimulation of immature fish with subsequent sampling of head kidney; **B)** *in vitro* stimulation of head kidney leukocytes of immature fish; 18 samples were used for RNA-Seq, 18 for ATAC-Seq and 18 for ChIP-Seq/µChIPmentation with antibodies to H3K4me3, H3K27ac and H3K27me3 histone marks.

### *In vitro* stimulation: leukocyte isolation and culture

Leukocytes were isolated from 12 fish. The entire head kidney was aseptically isolated and placed in a Petri dish with 40 ml of cell isolation media (500 ml of Leibovitz L-15 medium (L-15), 10 ml FBS (2 %), 0.02 % EDTA). Samples were then cut into small pieces and passed through a 100 µm nylon mesh with constant flow of cell isolation media. Leukocytes were separated by centrifugation of 40 ml of the cell suspension gently layered in a 50 ml tube containing 51 % Percoll (400 x g, 30 min, 4 °C, no brakes). The interface layer was collected by centrifugation (400 x g, 10 min, 4 °C) and washed three times with L-15 medium containing 0.1 FBS, keeping the pelleted leukocytes (Figure 7B).

To ensure enough cells were available, samples were pooled by pairs after cell counting and cell viability was evaluated by the trypan blue exclusion test, totalling 6 pools (each 18 x 10^6^ cells; Supplementary table 1). Then, each pool was divided into nine aliquots of 2 x 10^6^ cells (in 2 ml) that were dispensed into wells, for a total of 54 wells (6 pools x 9 aliquots): 18 stimulated with 20 µl of poly I:C solution; 18 with 20 µl of inactivated *V. anguillarum*; and the remaining 18 wells used as controls (Figure 7B). All leukocyte cultures were incubated for 24 h at 16 °C. Cells were then collected in 2 ml Eppendorf tubes and pelleted at 500 x g for 5 mins at room temperature. Eighteen pellets were resuspended in RNAlater and stored at -80 °C for RNA-Seq, while the other 36 were flash frozen in dry ice and stored at -80 °C for ATAC-Seq and µChIPmentation, respectively (Supplementary table 1).

### RNA isolation and sequencing

After sample thawing, total RNA was extracted and purified using the miRNeasy Kit (QIAGEN) with specific modifications for: i) frozen head kidney (> 20 mg), following a protocol for “Total RNA extraction for tissues”, and ii) frozen leukocytes (2 x 10^6^ cells) following a protocol for “Total RNA extraction for frozen cells” (Supplementary table 2). RNA integrity and quantity were evaluated in a Bioanalyzer (Bonsai Technologies, Madrid, Spain) and in a NanoDrop® ND-1000 spectrophotometer (NanoDrop® Technologies Inc., Wilmington, DE, USA). RNA integrity number (RIN) averaged 8.3 across all samples, always above 7.3. RNA samples were delivered to Novogene (UK) for library preparation using NEBNext Ultra Directional RNA Library Prep Kits for Illumina and sequenced using an Illumina NovaSeq S4 platform to generate 150 bp paired end reads.

### RNA-Seq data processing

RNA-Seq data was processed using nf-core/rnaseq 3.10.1 (Patel et al., 2023) run with default parameters, using the turbot Ensembl genome ASM1334776v1 (Martínez et al., 2021) as reference. In brief, the pipeline evaluated quality of raw reads using FASTQC (Andrews, 2010) and trimmed adapters and low-quality bases using Trim Galore! (Martin et al., 2011). Reads were then mapped using STAR (Dobin et al., 2013). Normalized transcript read counts were obtained using RSEM (Li and Dewey, 2011). After nf-core processing, the resulting count tables were filtered to remove genes with expression below 5 transcripts per million (TPM < 5) and represented in only one sample across all conditions.

### Differential gene expression and gene ontology analysis

Differentially expressed genes (DEGs) between stimulated and control samples were identified using the R/Bioconductor package DESeq2 v1.38.1 (Love et al., 2014). Genes with false discovery rate (FDR) adjusted p < 0.05 were considered DEGs. Functional enrichment of the DEG lists was performed using ShinyGO v0.77 (Ge et al., 2020). Gene ontology (GO) terms for Biological Process of each list were ranked by statistical significance (FDR-adjusted p < 0.05). All expressed genes across conditions were used as the background for GO analyses.

### ATAC-Seq: Library preparation and sequencing

Following a standard protocol for “Nuclei isolation for ATAC-Seq procedures” (Supplementary table 2), the frozen head kidney fragments (> 20 mg; *in vivo* assay) and cell pellets (2x10^6^ cells; *in vitro* assay) were thawed and resuspended in 1 ml of TST buffer. Each tissue fragment was cut into smaller pieces with a scalpel, mashed with the rubber back of a syringe, and filtered through a 40 µm cell strainer, while cell pellets were resuspended by gentle pipetting. The number and integrity of nuclei was assessed with a haemocytometer (minimum of ∼50,000 nuclei in a 16.5 µl suspension; ∼3,000 nuclei/µl) before carrying out the Tn5 transposase reaction with Illumina Tagment DNA TDE1 enzyme (37°C, 30 min, 1000 RPM, Illumina) following the standard “OmniATAC protocol” (Corces et al., 2017; Supplementary table 2). The resulting DNA was purified with a MinElute PCR purification kit (Qiagen), and DNA concentration assessed with a Qubit using the dsDNA HS kit (ThermoFisher Scientific). Library amplification (10-12 PCR cycles) was carried out using the NEBNext Ultra II DNA Library Prep Kit (New England Biolabs), with IDT for Illumina UD Indexes (96x, Plate A, Set 1, Illumina). Library size selection was performed to remove fragments below 180 bp and above 700 bp using AMPure XP beads (Beckman coulter). Finally, DNA fragment size distribution was assessed with the Bioanalyzer High Sensitivity DNA Assay kit (Agilent Technologies). ATAC-Seq libraries were delivered to Novogene (UK) to be sequenced on an Illumina NovaSeq S4 platform generating 150 bp paired end reads.

### ChIP-Seq and µChIPmentation: Library preparation and sequencing

The frozen head kidney fragments (> 20 mg) and leukocyte pellets (2x10^6^ cells) were thawed on ice. Following a standard “ChIP-seq” protocol (Supplementary table 2), the tissue fragments were transferred into a Douncer homogenizer containing a protease inhibitor cocktail (PIC, Roche, 1 tablet in 50 ml of PBS) solution immersed in ice and homogenized using pestles A and B (from less to more plunger adjustment). The leukocyte pellets were resuspended in the PBS and PIC solution. The amount and quality of nuclei was assessed with a haemocytometer and trypan blue staining (> 10 million cells for head kidney; > 100,000 for leukocyte cultures). Due to the low number of nuclei recovered from the leukocyte cultures, a µChIPmentation protocol (Diagenode) was used, following a modified protocol (Supplementary table 2).

In both cases, chromatin crosslinking was done using a 1 % formaldehyde solution followed by quenching with glycine (0.125 M). Nuclei were pelleted and resuspended in complete sonication buffer, while leukocyte nuclei were resuspended in Hanks’ Balanced Salt Solution (HBSS, Thermo Fisher Scientific) - tL1 buffer (Supplementary table 2). The chromatin was sheared using a Covaris S2 focused ultrasonicator with the following parameters: 2 % duty cycle, intensity 3, with 200 cycles per burst, at 4 °C for 8 min and 6 min for head kidney tissue and leukocytes, respectively.

The immunoprecipitation was performed using Diagenode antibodies for three marks: H3K4me3 (marking active promoter regions; cat. No. C15410003; 1.3 µg/µl), H3K27ac (marking active enhancer and promoter regions; cat. No. C15410196; 2.8 µg/µl) and H3K27me3 (marking Polycomb repressed regions; cat. No. C15410195; 1.1 µg/µl). For head kidney samples, antibodies were coupled with pre-washed protein A and protein G beads, and the tubes left under rotation overnight (∼16 h) at 4 °C, following the standard ChIP-Seq protocol. After washing the beads and decrosslinking, the samples were purified using the MinElute PCR purification kit (Qiagen) and later quantified with DNA HS Qubit (ThermoFisher Scientific). Immunoprecipitated chromatin was stored at -20 °C until library preparation, using a Microplex v3 kit (Diagenode).

For the leukocyte samples, the µChipmentation kit for histones (Diagenode) was used for chromatin immunoprecipitation and ChIP-Seq library preparation (Supplementary table 2).

Before sequencing, the quantity and quality of purified libraries was assessed using the Qubit DNA HS kit (ThermoFisher Scientific) and the High Sensitivity DNA Assay kit (Agilent Technologies), respectively. A minimum of 60 % of the chromatin was required to have a size distribution between 200-700 bp (centered around 350-400 bp). ChIP-Seq libraries were delivered to Novogene (UK) for sequencing on an Illumina NovaSeq S4 platform generating 150 bp paired-end reads.

### ATAC-Seq and ChIP-Seq data processing

ATAC-Seq and ChIP-Seq data were processed using the nf-core/atacseq v1.2.2. and nf-core/chipseq v1.2.2 pipelines (Ewels et al., 2023), respectively, run with the narrow_peak option for ATAC-Seq and H3K4me3 and H3K27ac ChIP-Seq datasets, and with the broad_peak option for H3K27me3. The other parameters were kept by default. Quality assessment of the reads was carried out with FASTQC (Andrews, 2010), and adapters and low-quality bases trimmed with Trim Galore! (Martin et al., 2011). Reads were mapped to the turbot genome using BWA (Li and Durbin, 2009). Further filtering was done with SAMtools (Danecek et al., 2021), BAMtools (Barnett et al., 2011) and Pysam (Danecek et al., 2021). Genome-wide immunoprecipitation (IP) enrichment relative to controls was done with deepTools (Ramírez et al., 2014) and broad/narrow peaks were called using MACS2 (Zhang et al., 2008). Once nf-core was finished, suboptimal replicates with very low peak numbers were excluded after visualizing their bigwig files on the Integrative Genomics Viewer (IGV; Robinson et al., 2023).

### ChIP-Seq and µChIPmentation blacklist

To improve the signal-to-noise ratio of the ChIP-Seq and µChIPmentation data, a blacklist consisting of high signal and low mappability regions was constructed using ChIP-Seq and µChIPmentation inputs, including 21 control ChIPseq turbot samples (ENA accession PRJEB57784). The mappability of the turbot genome for read lengths of 100 bp and 150 bp (k-mers 100 and 150) was quantified using the umap software package (Karimzadeh et al., 2018). The generated mappability files were fed into the ENCODE blacklist software (Amemiya et al., 2019) to generate the blacklist. Alignment and peak files derived from the nf-core pipeline were filtered with BAMtools to remove reads and peaks located in the blacklist regions.

### Differential histone modification regions and differentially accessible regions

Differential histone modification regions (DHMRs) and differentially accessible regions (DARs; adjusted p < 0.05) between stimulated samples and controls were identified using DiffBind (Stark and Brown, 2011) with default settings.

### Integration of ChIP-Seq and ATAC-Seq data with RNA-Seq data

For each condition tested, the promoters of DEGs overlapping with regions tagged as DARs and/or DHMRs were identified. When applicable, a hypergeometric test was performed to check the significance of overlapping between each pair (p < 0.05).

### Chromatin state inferences

Genome-wide chromatin states for each condition were predicted using ChromHMM (Ernst and Kellis, 2012) integrating the ChIP-Seq data (µChIPmentation for *in vitro* samples) for the three histone marks (H3K4me3, H3K27ac and H3K27me3) and ATAC-Seq data. Chromatin state prediction was performed by testing ChromHMM models including from 8 to 15 states, keeping the one that returned the most biologically relevant chromatin states, for head kidney and leukocyte data separately (Pan et al., 2021; Baranasic et al., 2022; Vu and Enst, 2022). The genome-wide distribution of resulting chromatin states was visualized on IGV. Regions annotated as enhancer-related states by ChromHMM were retrieved, and each stimulation dataset was compared against its respective control to identify potential enhancer-related regions. Then, each list of regions including potential enhancers annotated as “intergenic” or “intron” were kept as differential enhancer-state regions for further analysis.

### Transcription factor motif analysis

Enriched transcription factor (TF) binding motifs (TFBM) included in the HOMER software (Heinz et al., 2010) were identified using the *findMotifsGenome.pl* function (settings: -size given -mask -mset vertebrates) in the different lists of promoter-associated DHMRs/DARs and putative enhancers for each condition. Random genomic regions with GC-content matching each input genomic list were used as the background for automatic motif analysis by HOMER. Only TFBMs with p < 0.01 and percentage of target sequences > 5 % were considered enriched.

GO analysis of the TFs predicted to bind enriched TFBMs was performed with Metascape (Zhou et al., 2019) using zebrafish *Danio rerio* as the reference. Finally, a selection of TF genes that were DEGs and contained promoter regions overlapping DARs or DHMRs were explored with IGV (Robinson et al., 2023).

## DATA ACCESS

All raw RNA-Seq, ATAC-Seq and ChIP-Seq datasets can be accessed through the ENA repository under accession numbers PRJEB47933, PRJEB47934 and PRJEB57784, respectively. Detailed metadata for the samples and prepared libraries are available in Supplementary tables 1 and 2, respectively. Detailed experimental protocols are publicly available in the FAANG repository (data.faang.org) and following the URLs facilitated in Supplementary tables 1 and 2.

## Supporting information

Supplementary material descriptions

## ACKNOWLEDGMENTS

We acknowledge the technical support and informatic resources provided by the Centro de Supercomputación de Galicia (CESGA). We acknowledge Professor Carolina Tafalla (INIA-CSIC) for the support given in the revision of this article.

## AUTHOR CONTRIBUTIONS

**OA:** Methodology, Software, Formal analysis, Investigation, Data Curation, Visualization, Writing original draft, Writing – review & editing; **BGP:** Methodology, Investigation, Resources, Writing review & editing; **PRV:** Investigation, Resources, Writing – review & editing; **ABH:** Software, Formal Analysis, Data Curation, Visualization, Writing – review & editing; **JL**: Investigation, Resources, Writing – review & editing; **PD:** Software, Formal Analysis, Writing – review & editing; **DPM:** Software, Formal Analysis, Writing – review & editing; **PB**: Resources, Writing – review & editing; **DM**: Funding Acquisition, Methodology, Supervision, Resources, Writing – review & editing; **CB**: Conceptualization, Formal Analysis, Resources, Writing – review & editing, Project Administration, Supervision; **PM**: Conceptualization, Methodology, Formal Analysis, Resources, Writing – review & editing, Project Administration, Supervision.

