## Supplementary Figures 1 to 5 for "Multiomics uncovers the epigenomic and transcriptomic response to viral and bacterial stimulation in turbot"

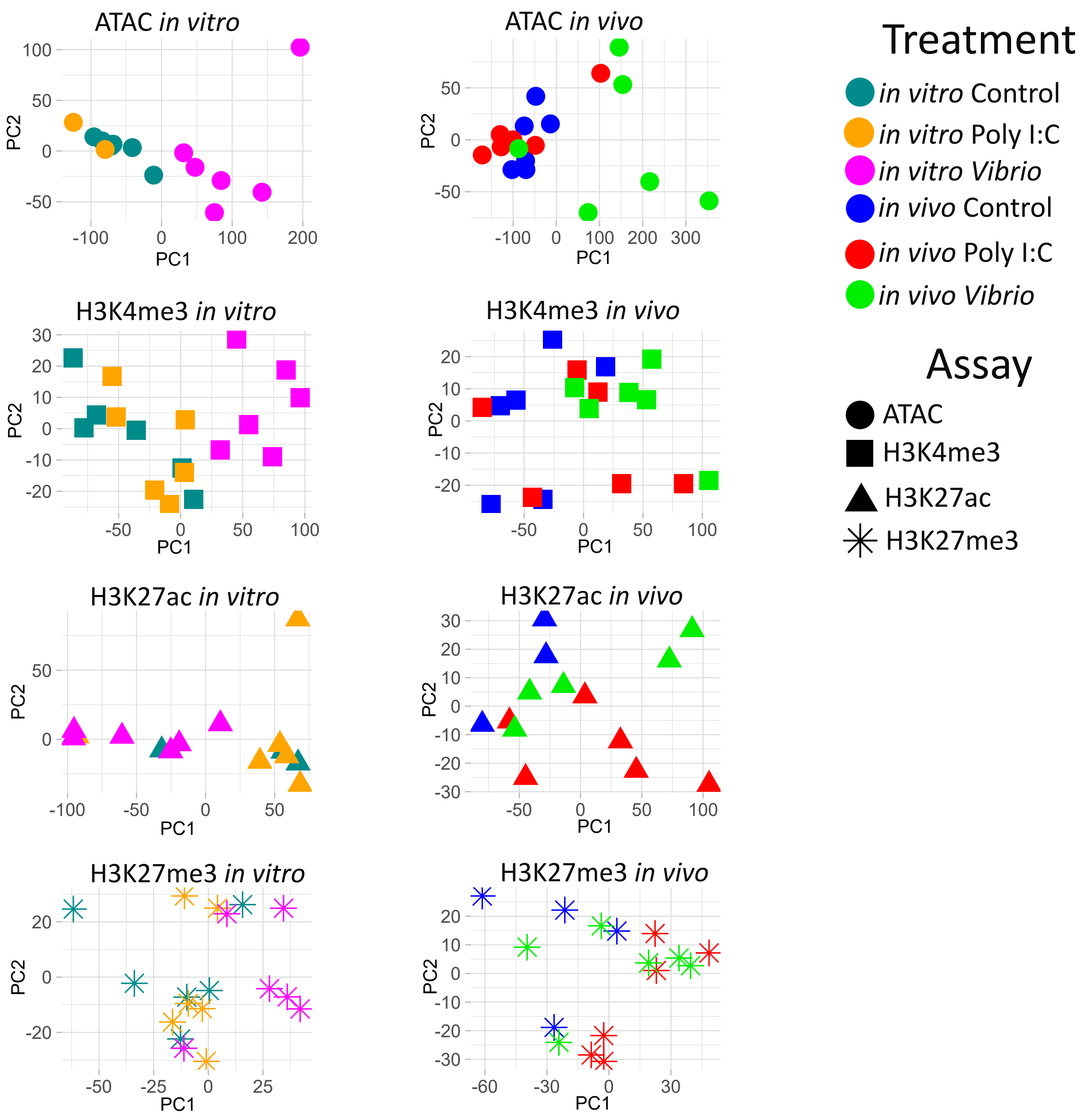


**Supplementary Figure 1.** Separate PCAs for each assay and stimulation of the *in vivo* and *in vitro* epigenomic response to stimulation with poly I:C and *Vibrio* for the open chromatin ATAC-Seq and histone ChIP-Seq (H3K4me3, H3K27ac, H3K27me3) samples.

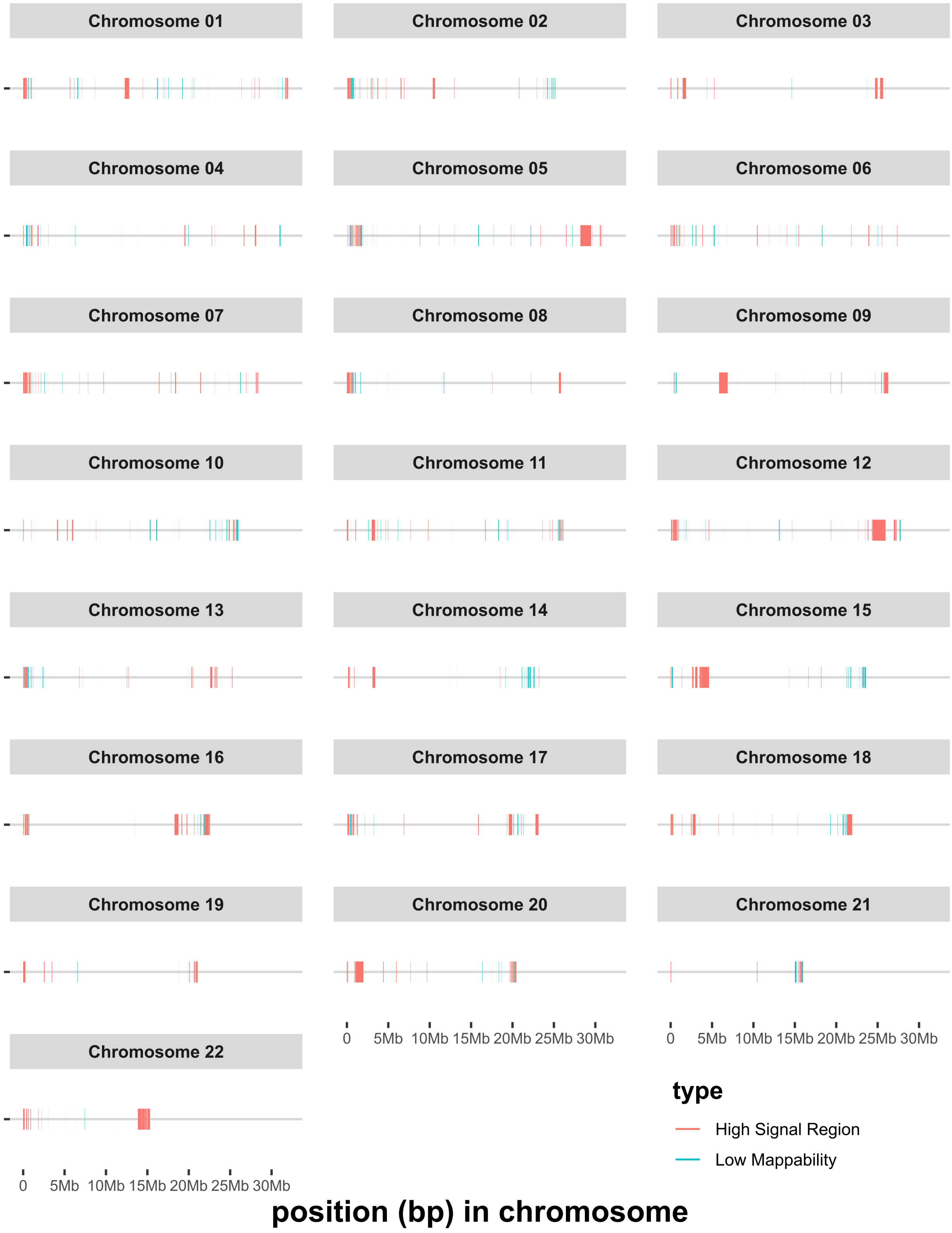


**Supplementary Figure 2.** Genomic distribution of the ChIP-Seq high signal regions (pink) and low mappability regions (blue) of the turbot genome included in the blacklist for each of the 22 chromosomes.

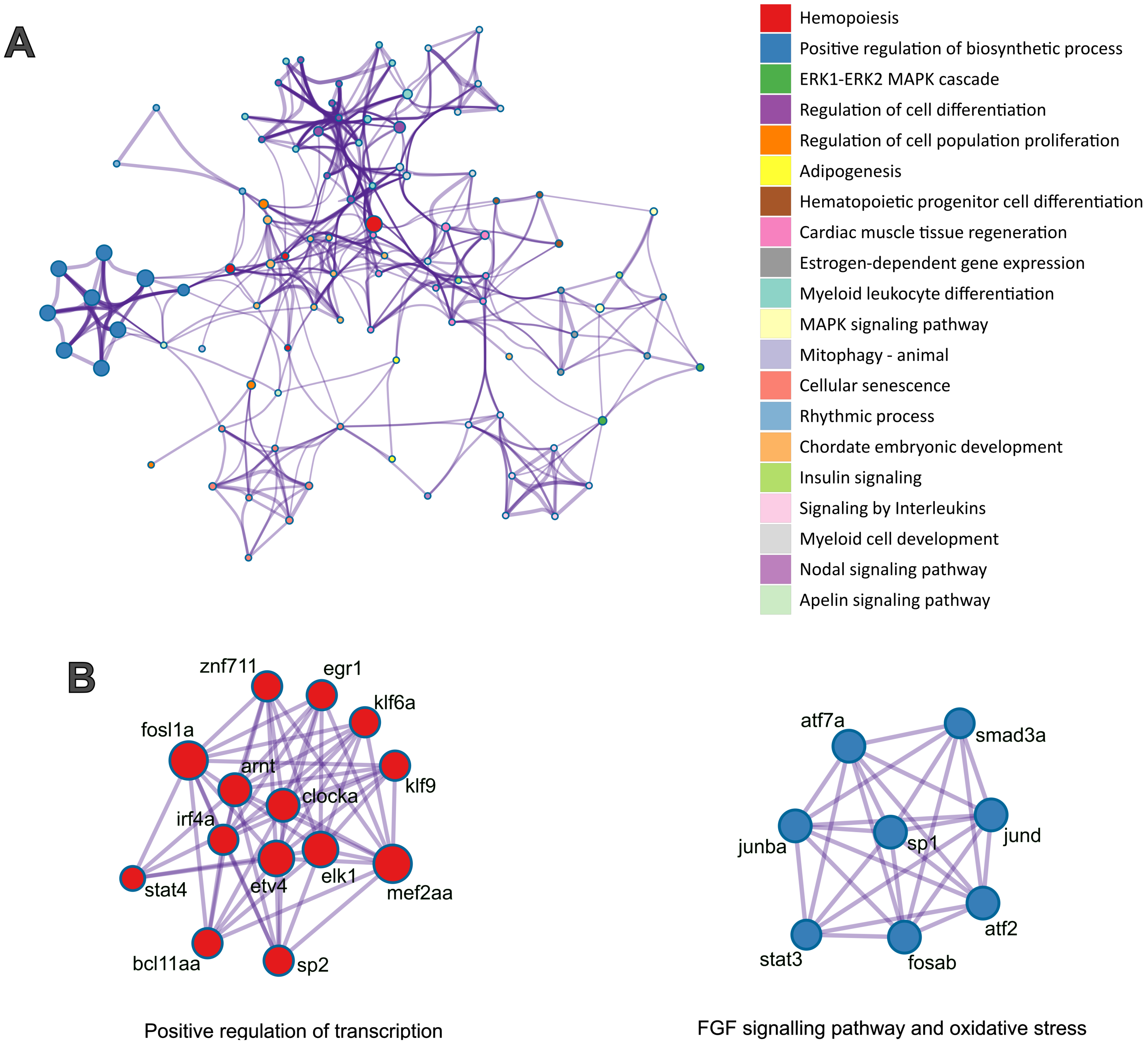


**Supplementary Figure 3.** Functional interconection of active TFs detected in response to teh different stimulants: A) Ontology enrichment clusters of upregulated TFs showing promoter DARs/DHMRs with TFBMs enriched in promoter or enhancer DARs/DHMRs of DEGs detected in Vibrio in vivo stimulation. B) Protein-protein interaction MCODE network of upregulated TFs induced by Vibrio in vivo stimulation. Each cluster is coloured with the most statistically significant term among terms that cluster together. The size of each term is given by -log10 p, and the stronger the similarity among nearby terms, the thicker the edges between them

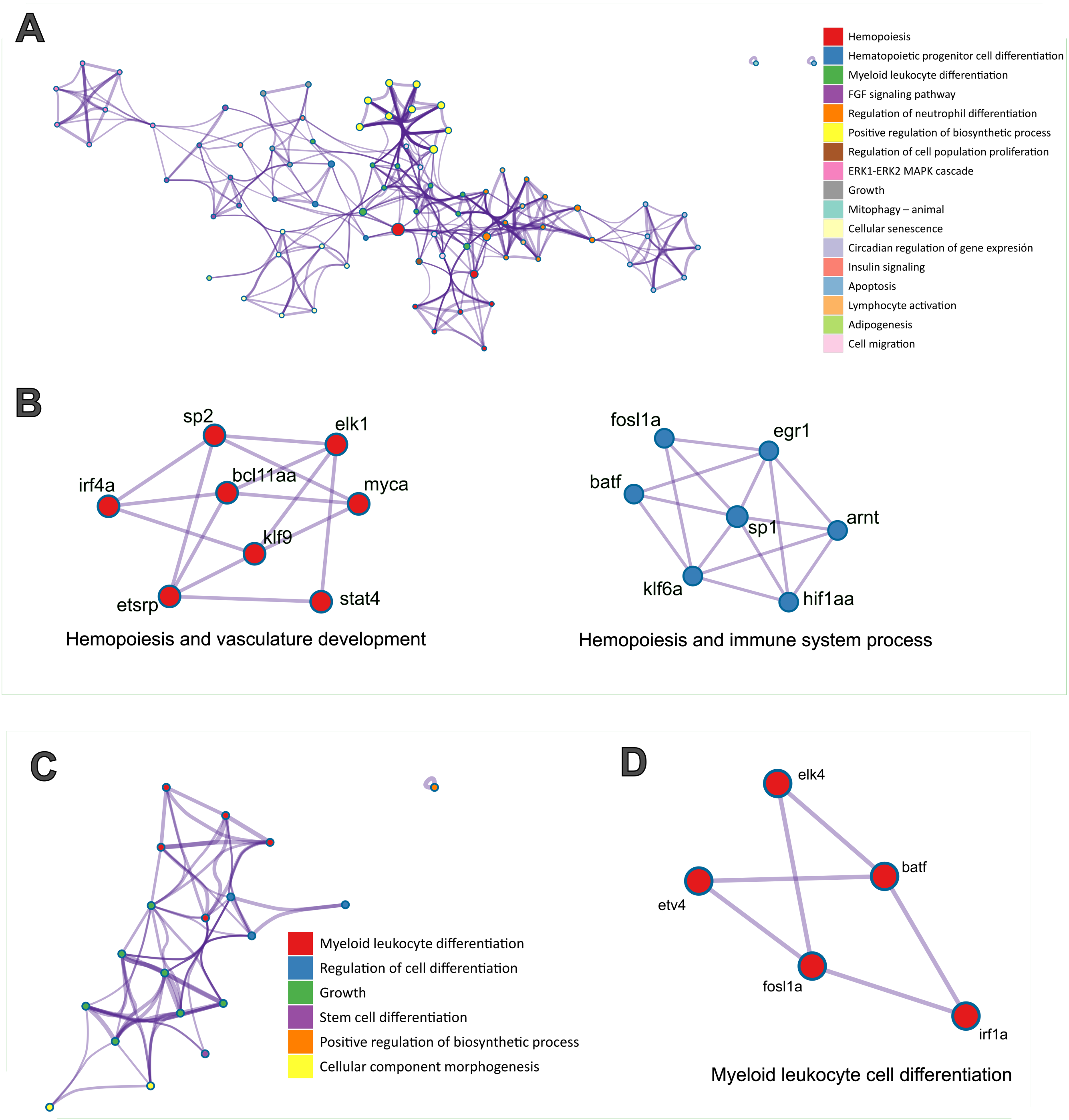


**Supplementary Figure 4.** Functional interconection of active TFs detected in response to teh different stimulants: A) and C) Ontology enrichment clusters of upregulated TFs showing promoter DARs/DHMRs with TFBMs enriched in promoter or enhancer DARs/DHMRs of DEGs detected in A) *Vibrio in vitro* and C) poly I:C *in vivo* stimulation. B) and D) Protein-protein interaction MCODE network of upregulated TFs induced by B) *Vibrio in vitro* and D) poly I:C *in vivo* stimulation. Each cluster is coloured with the most statistically significant term among terms that cluster together. The size of each term is given by -log10 p, and the stronger the similarity among nearby terms, the thicker the edges between them.

### arnt (HIF1B)
### Chromosome 1: 19,174,451-19,183,553

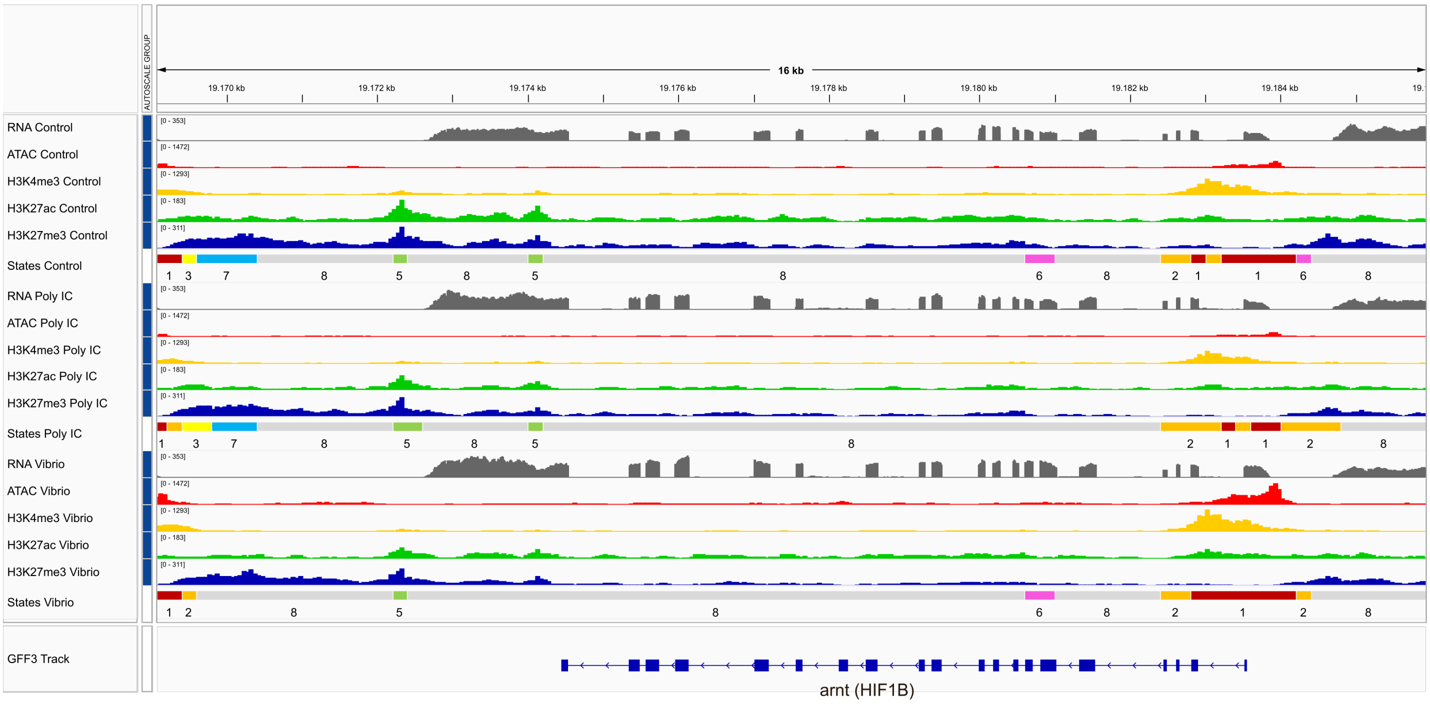


### irf8
### Chromosome 5: 14,893,961-14,898,170

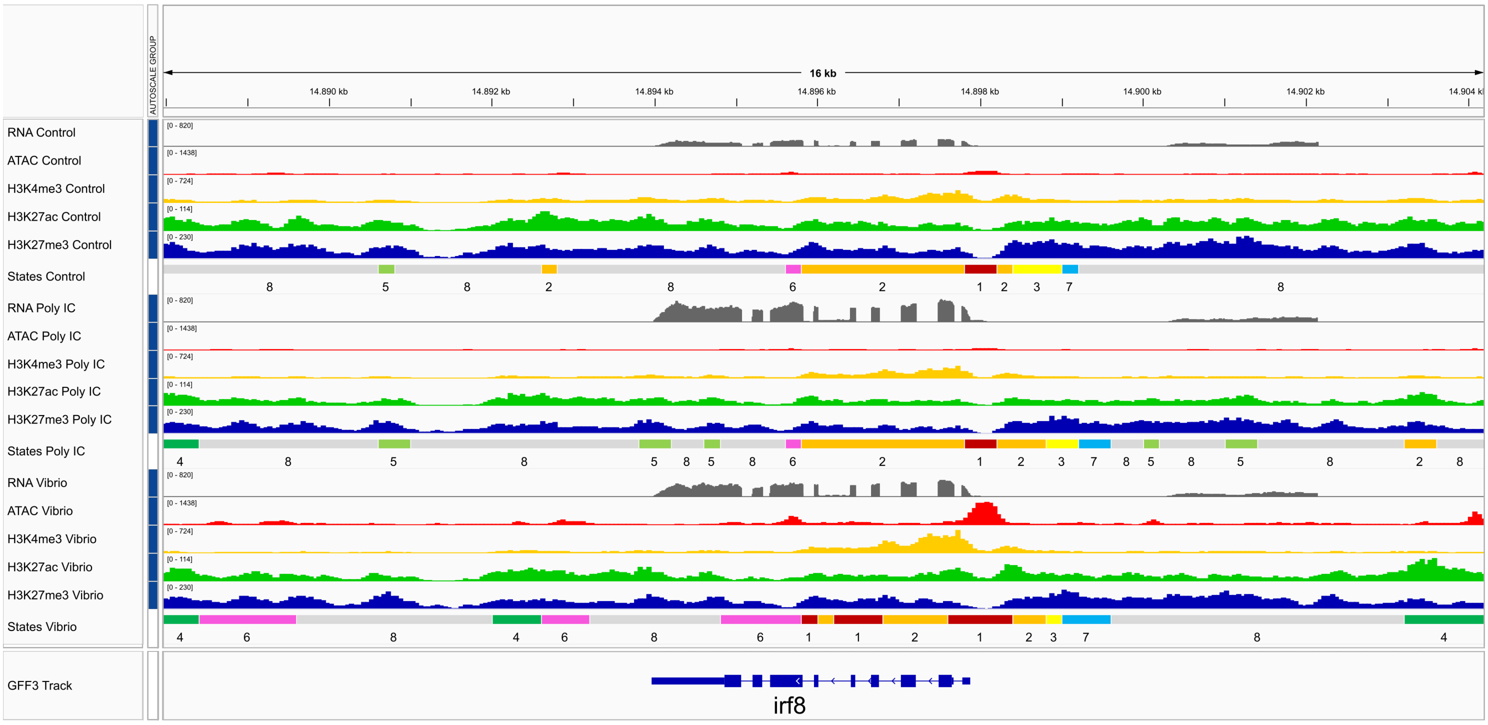


| | | | | | | | |
|---|---|---|---|---|---|---|---|
| 1 | Strongly active promoter / transcript | 3 | Bivalent / poised TSS | 5 | Bivalent / poised enhancer | 7 | Repressed Polycomb |
| 2 | Flanking active TSS without ATAC | 4 | Strong active enhancer | 6 | ATAC island | 8 | Quiescent / Low |

**Supplementary Figure 5A.** IGV screenshots of the chromatin structure and RNA-Seq, ATAC-Seq and ChIP-Seq tracks of different transcription factor coding genes, selected due to them i) being differentially expressed (DE) in at least one of the conditions; ii) having differentially accesible regions (DAR) or differential histone modification regions (DHMR) in at least one of the conditions and iii) being among the TFBM enriched in response to at least one of the conditions. Each chromatin state model annotation is located in the table bellow the IGV screenshots. *arnt* was DE and showed DAR and H3K4me3-DAR in its promoter in *Vibrio in vitro*, as well as being among the TFBM enriched in response to both *Vibrio in vitro* and *in vivo*. *irf8* was DE in both *Vibrio* and Poly IC *in vitro*, but only showed DAR in its promoter in *Vibrio in vitro*, as well as being among the enriched TFBM in response to *Vibrio in vitro*, *in vivo* and Poly IC *in vivo*.

bhlhe40a
Chromosome 6: 14,702,766-14,705,188

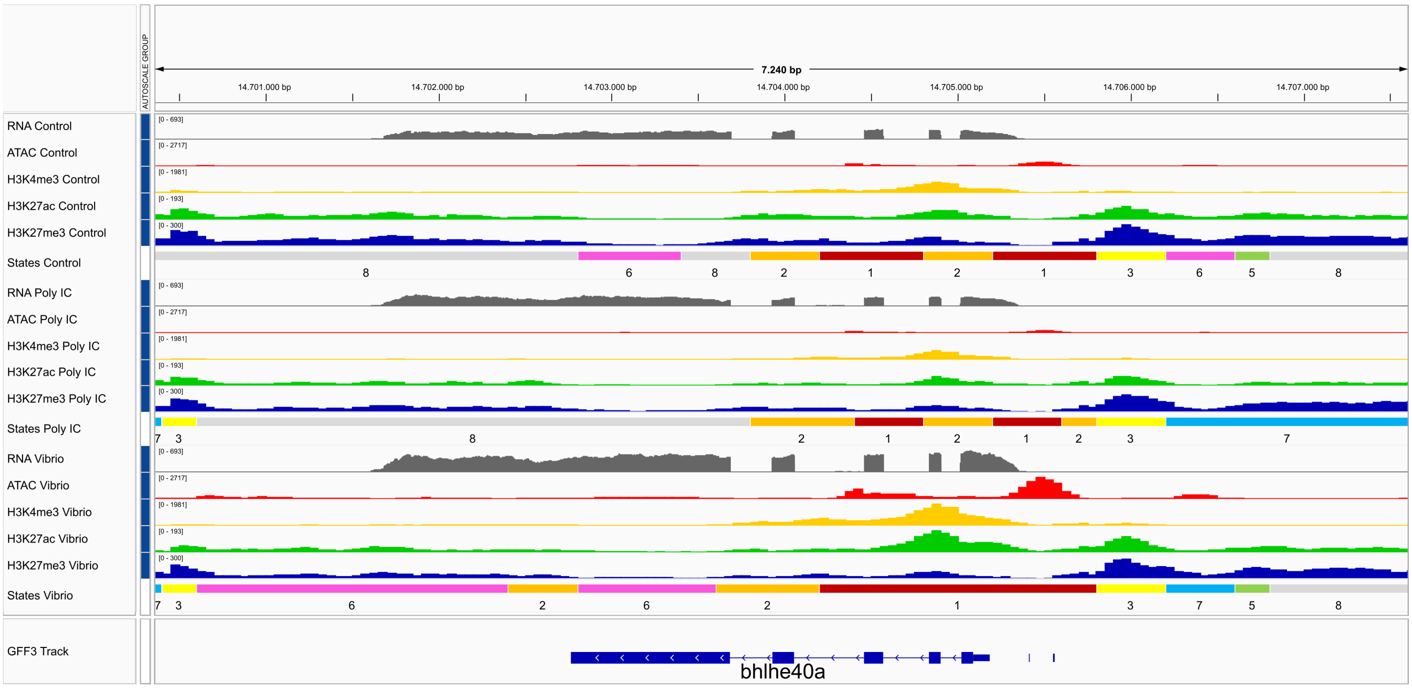


bhlhe40b
Chromosome 6: 21,238,776-21,240,855

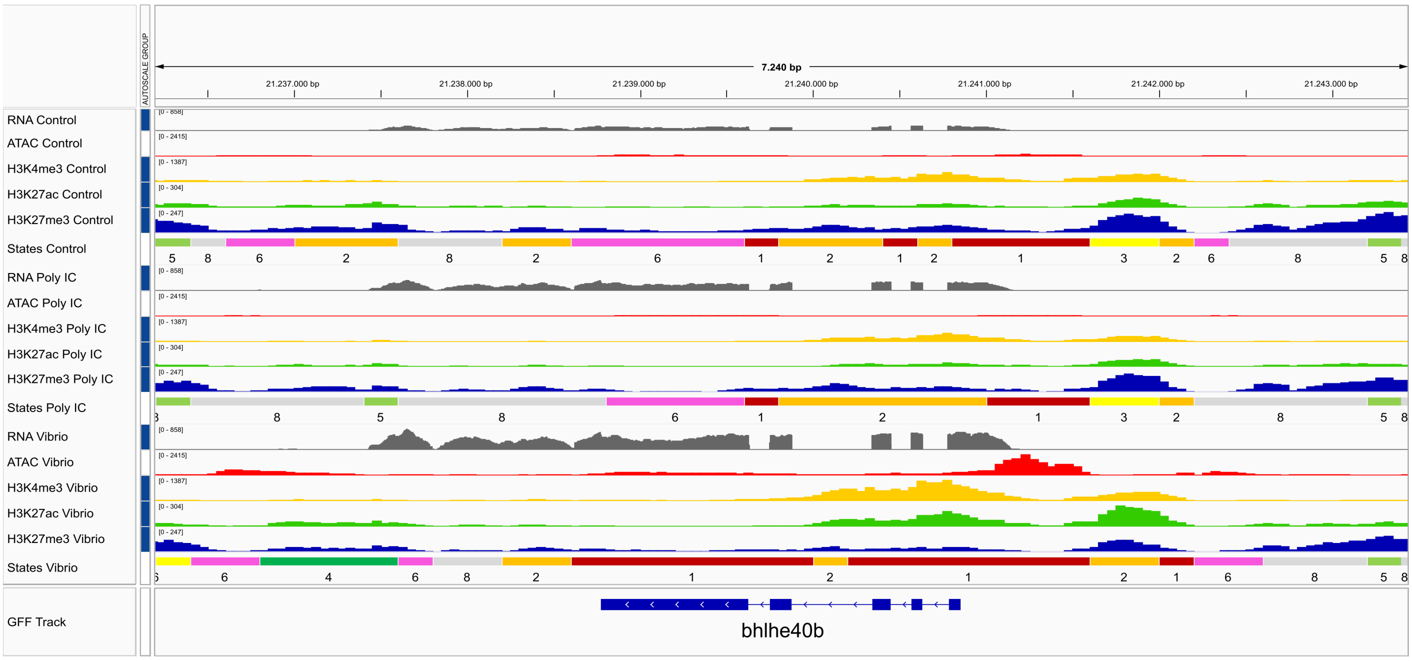


| | | | | | | | |
|---|---|---|---|---|---|---|---|
| 1 | Strongly active promoter / transcript | 3 | Bivalent / poised TSS | 5 | Bivalent / poised enhancer | 7 | Repressed Polycomb |
| 2 | Flanking active TSS without ATAC | 4 | Strong active enhancer | 6 | ATAC island | 8 | Quiescent / Low |

**Supplementary Figure 5B.** (Cont.) *bhlhe40a* and *bhlhe40b* were DE and showed DAR in its promoter in *Vibrio in vitro*, as well as being among the enriched TFBM in response to *Vibrio in vitro*, *in vivo*, and Poly IC *in vivo*.

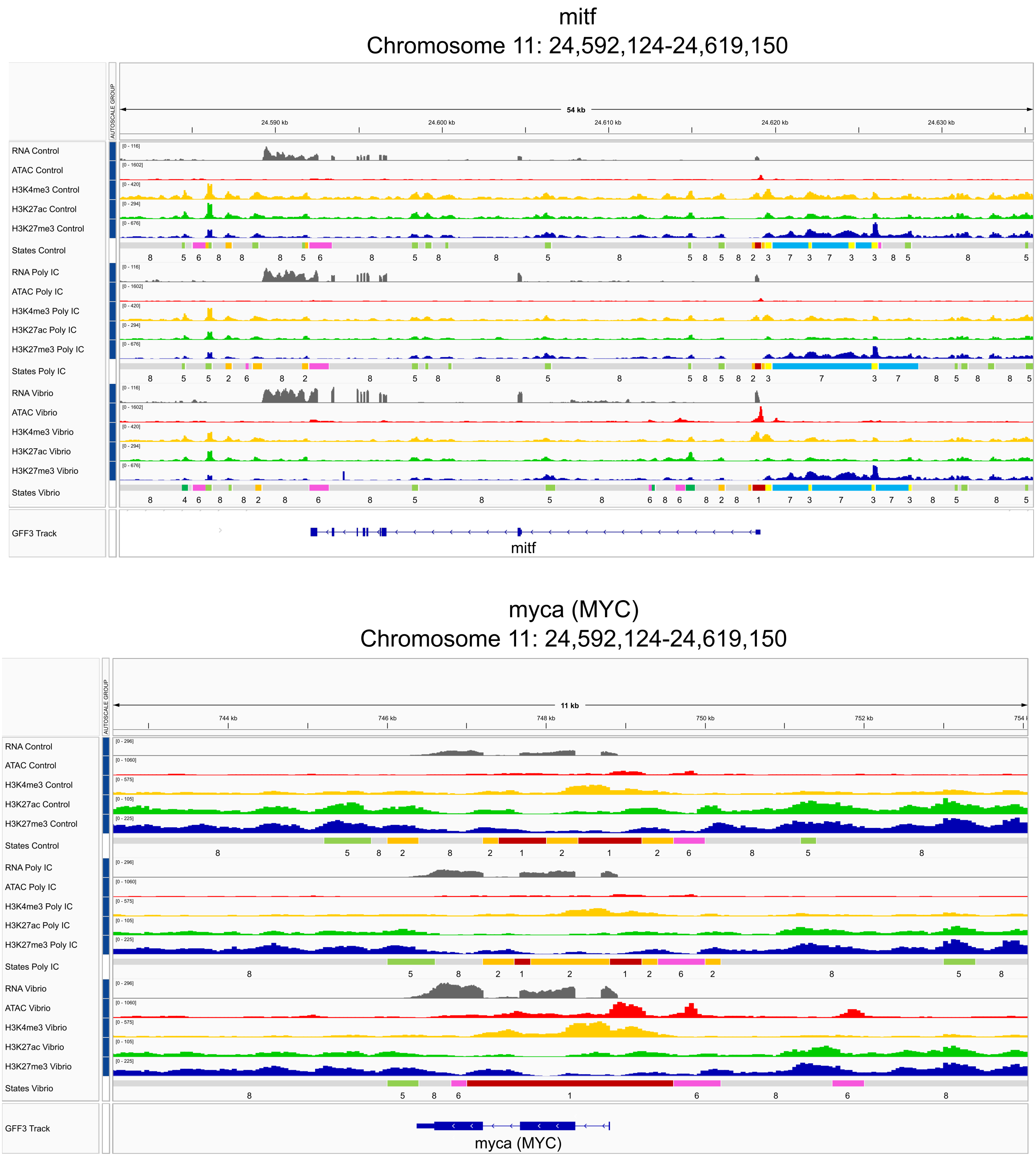


| | | | | | | | |
|---|---|---|---|---|---|---|---|
| 1 | Strongly active promoter / transcript | 3 | Bivalent / poised TSS | 5 | Bivalent / poised enhancer | 7 | Repressed Polycomb |
| 2 | Flanking active TSS without ATAC | 4 | Strong active enhancer | 6 | ATAC island | 8 | Quiescent / Low |

**Supplementary Figure 5C.** (Cont.) *mitf* and *myca* were DE and showed DAR in its promoter in *Vibrio in vitro*, as well as being among the enriched TFBM in response to *Vibrio in vitro*, *in vivo*, and Poly IC *in vivo*.

#### irf3
#### Chromosome 18: 12,159,911-12,165,967

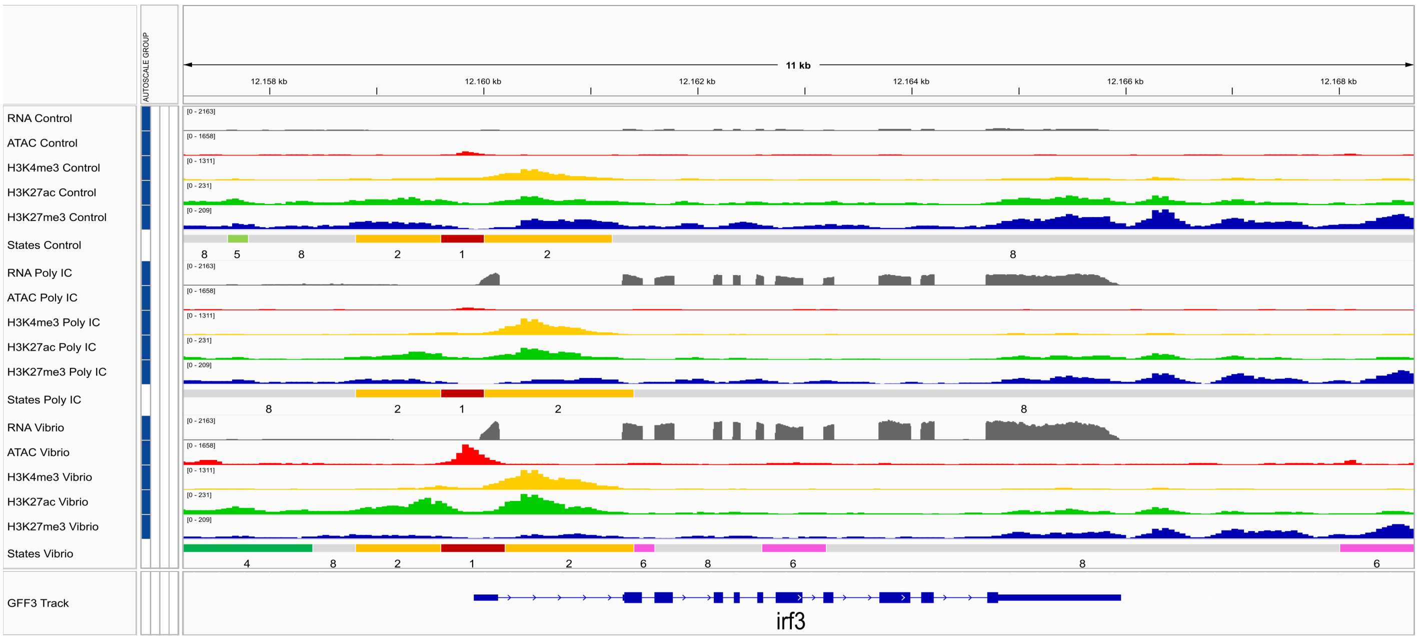


| | | | | | | | |
|---|---|---|---|---|---|---|---|
| 1 | Strongly active promoter / transcript | 3 | Bivalent / poised TSS | 5 | Bivalent / poised enhancer | 7 | Repressed Polycomb |
| 2 | Flanking active TSS without ATAC | 4 | Strong active enhancer | 6 | ATAC island | 8 | Quiescent / Low |

#### zeb2
#### Chromosome 1: 10,034,126-10,120,005

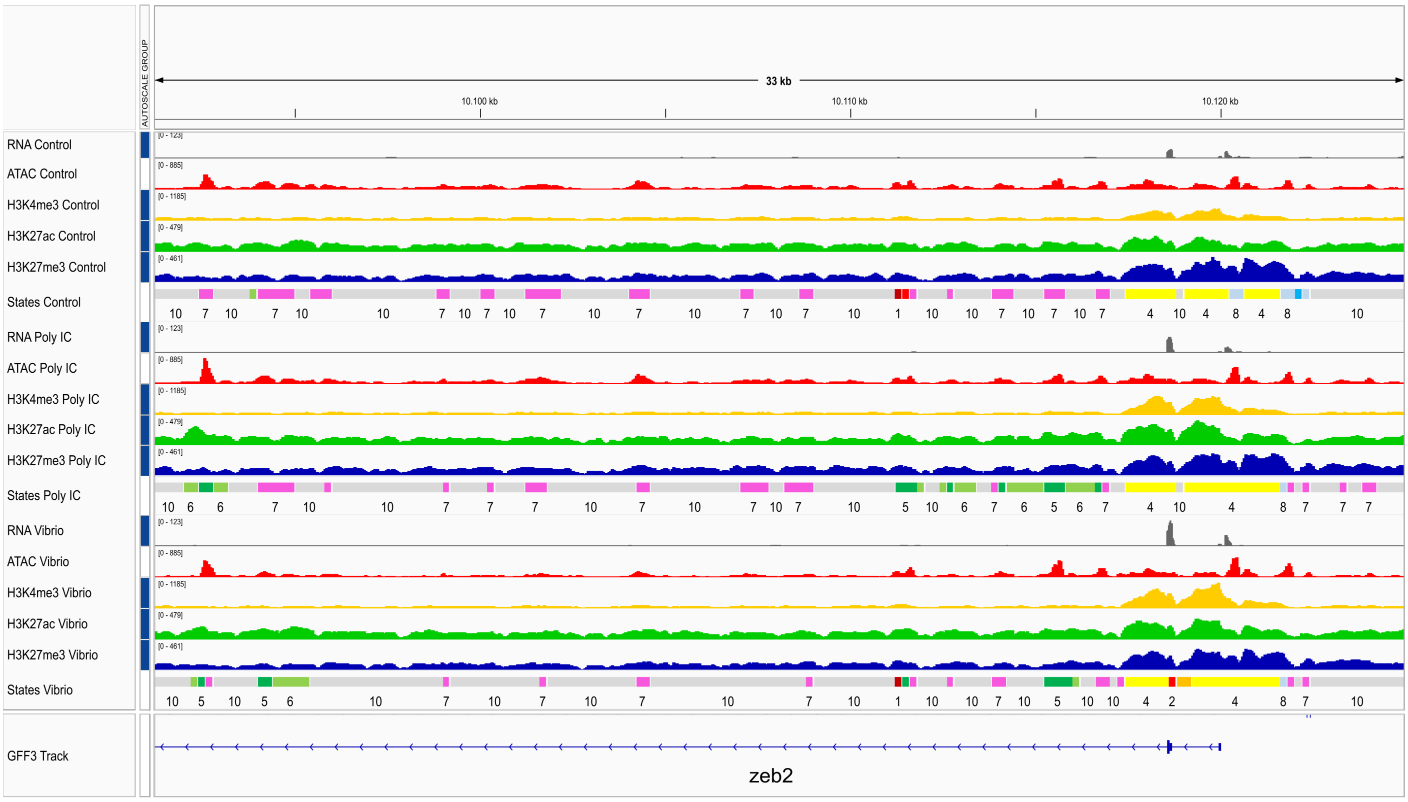


| | | | | | | | | | |
|---|---|---|---|---|---|---|---|---|---|
| 1 | Strongly active promoter / transcript | 3 | Flanking active TSS without ATAC | 5 | Strong active enhancer | 7 | ATAC island | 9 | Repressed Polycomb |
| 2 | Weak active promoter / transcript | 4 | Bivalent/poised TSS | 6 | Weak active enhancer | 8 | Weak Repressed Polycomb | 10 | Quiescent / Low |

**Supplementary Figure 5D.** (Cont.) *irf3* was DE in both *Vibrio* and Poly IC in vitro, but showed DAR in its promoter only in *Vibrio in vitro*, as well as being among the enriched TFBM in response to *Vibrio in vitro*, *in vivo*, and Poly IC *in vivo*. *zeb2* was DE and showed H3K4me3-DHMR in its promoter in *Vibrio in vivo*, as well as being among the enriched TFBM in response to *Vibrio in vivo*, *in vitro*, and Poly IC *in vivo*.

### egr1
#### Chromosome 13: 14,094,894-14,099,153

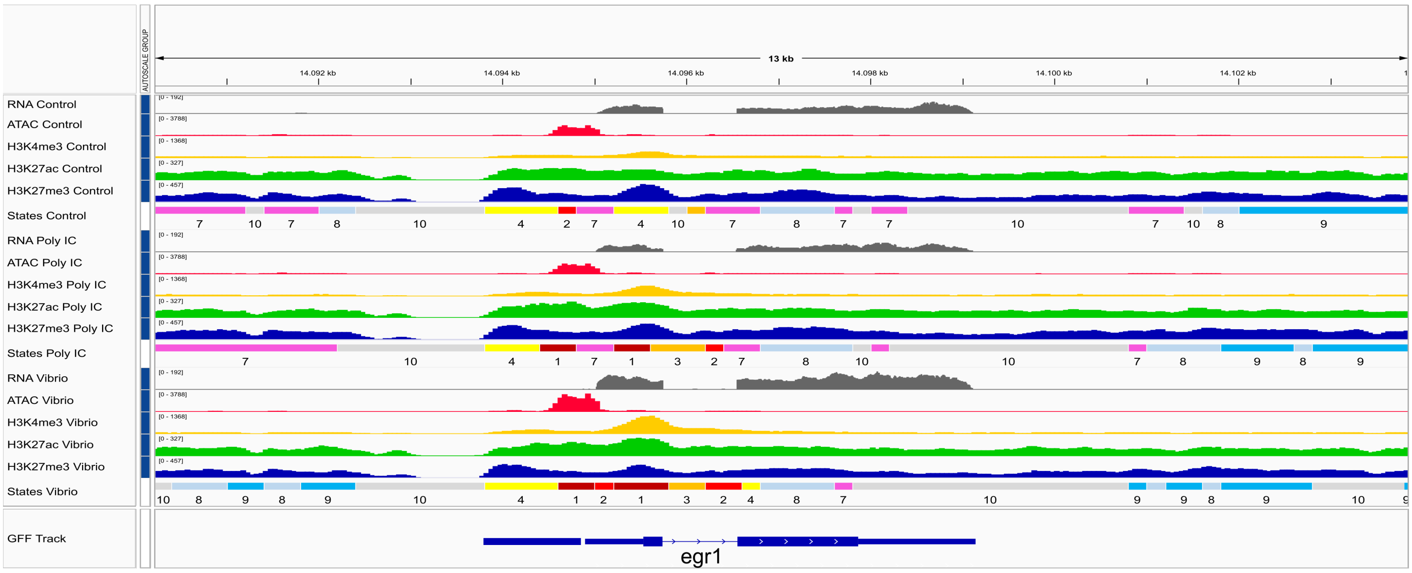


### stat4
#### Chromosome 14: 389,657-402,701

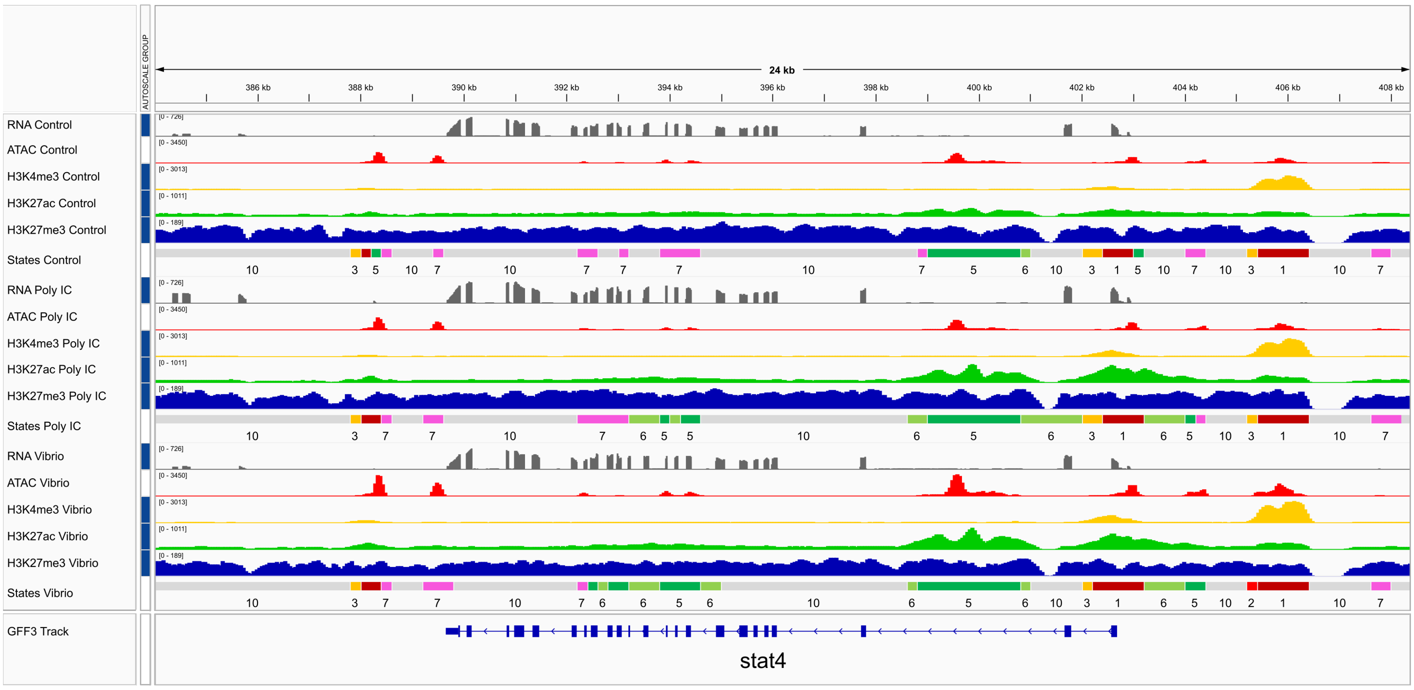


| | | | | | | | | | |
|---|---|---|---|---|---|---|---|---|---|
| 1 | Strongly active promoter / transcript | 3 | Flanking active TSS without ATAC | 5 | Strong active enhancer | 7 | ATAC island | 9 | Repressed Polycomb |
| 2 | Weak active promoter / transcript | 4 | Bivalent/poised TSS | 6 | Weak active enhancer | 8 | Weak Repressed Polycomb | 10 | Quiescent / Low |

**Supplementary Figure 5E.** (Cont.) *egr1* was DE and showed DAR in its promoter in *Vibrio in vivo*, as well as being among the enriched TFBMs in response to both *Vibrio in vivo* and *in vitro*. *stat4* was DE in both *Vibrio* and Poly IC *in vivo*, but showed DAR in its promoter only in *Vibrio in vivo*, as well as being among the enriched TFBM in response to *Vibrio in vivo*, *in vitro*, and Poly IC *in vivo*.

bcl11a
Chromosome 15: 16,712,975-16,756,991

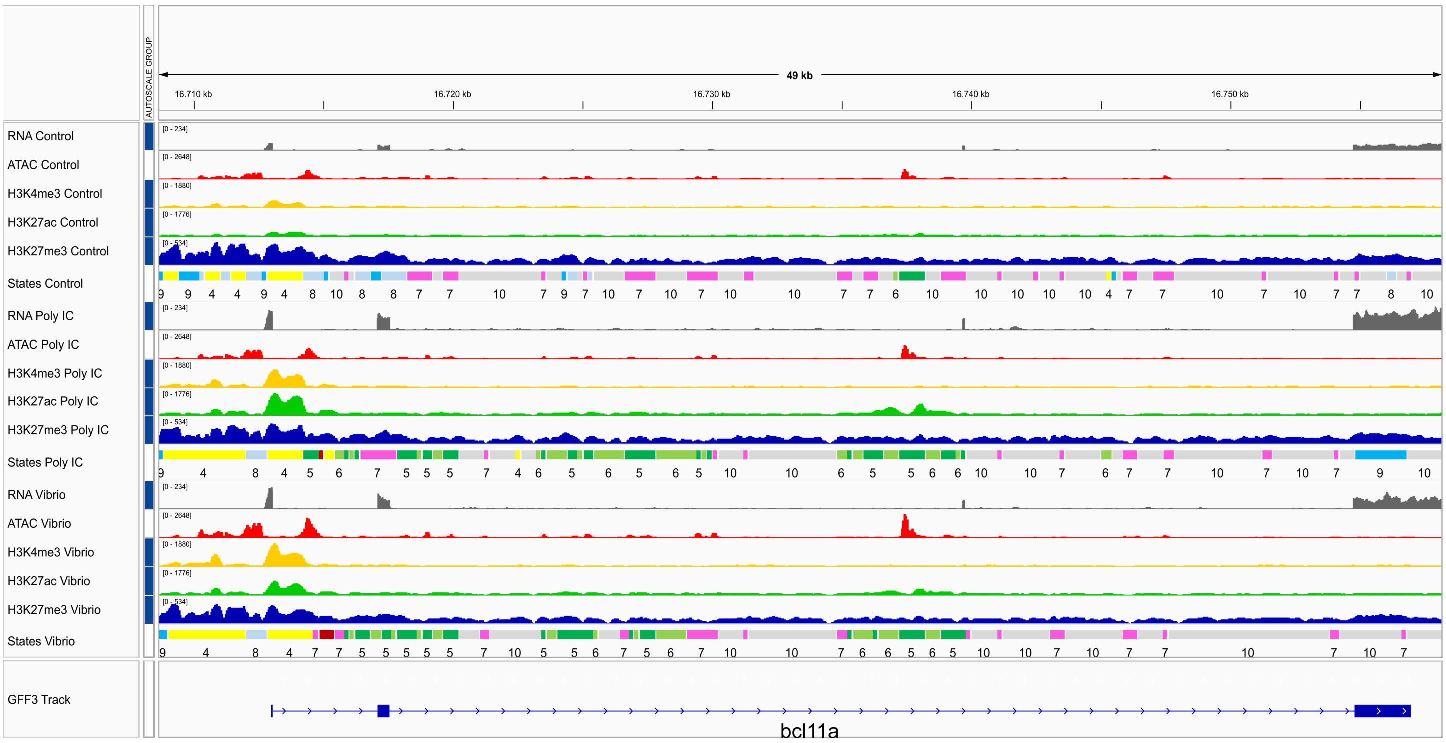


meis1b
Chromosome 15: 18,783,030-18,864,551

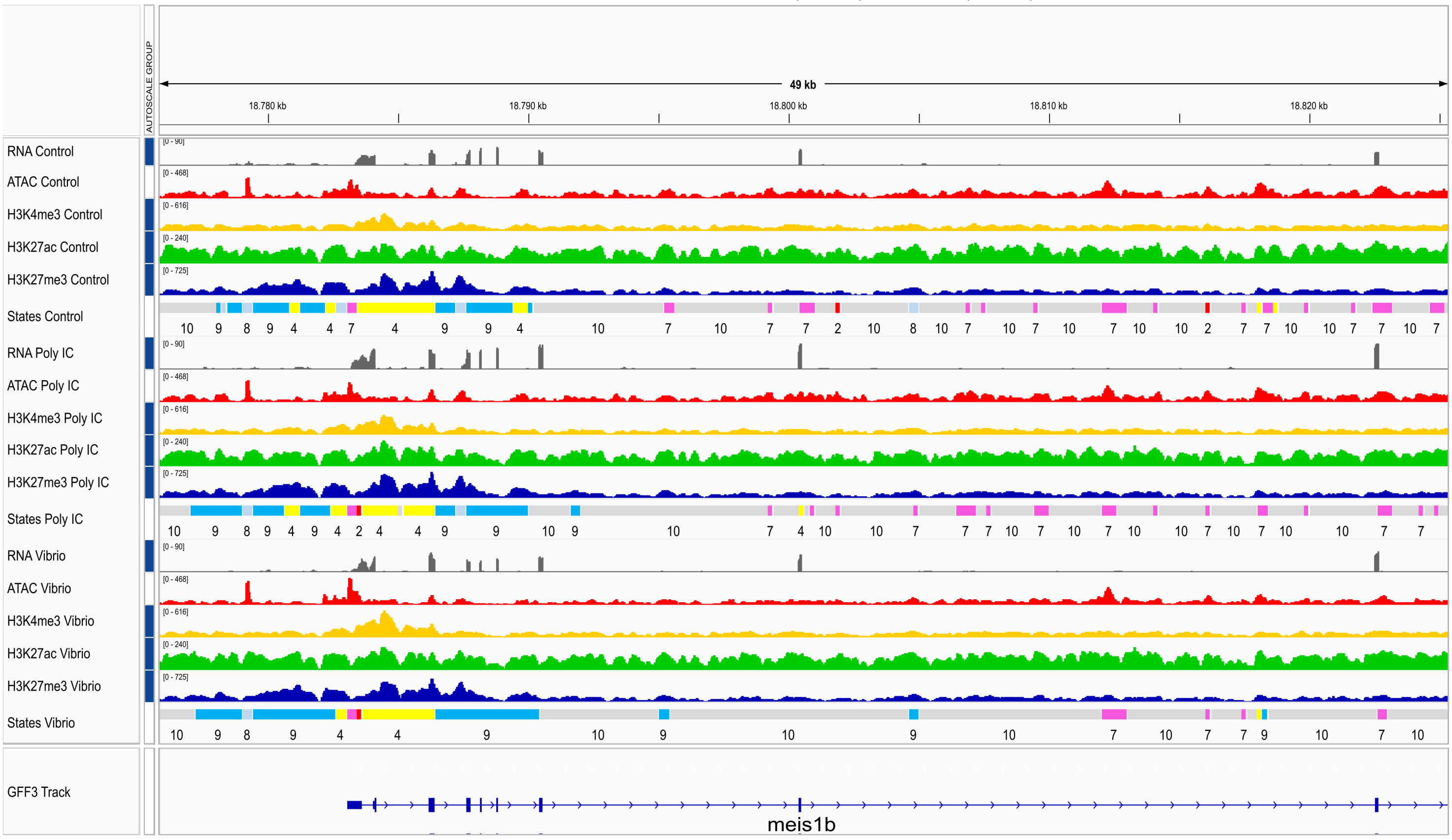


| | | | | | | | | | |
|---|---|---|---|---|---|---|---|---|---|
| 1 | Strongly active promoter / transcript | 3 | Flanking active TSS without ATAC | 5 | Strong active enhancer | 7 | ATAC island | 9 | Repressed Polycomb |
| 2 | Weak active promoter / transcript | 4 | Bivalent/poised TSS | 6 | Weak active enhancer | 8 | Weak Repressed Polycomb | 10 | Quiescent / Low |

**Supplementary Figure 5F.** (Cont.) *bcl11a* was DE in both Vibrio and Poly IC *in vivo*, but showed DAR in its promoter only in *Vibrio in vivo*, as well as being among the enriched TFBM is response to both *Vibrio in vivo* and *in vitro*. *meis1b* was DE and showed DAR its promoter in *Vibrio in vivo*, as well as being among the enriched TFBMs in response to both *Vibrio in vivo* and *in vitro*.
