## Supplementary material descriptions for "Multiomics uncovers the epigenomic and transcriptomic response to viral and bacterial stimulation in turbot"

**Supplementary Figure 1.** Separate PCAs for each assay and stimulation of the *in vivo* and *in vitro* epigenomic response to stimulation with poly I:C and Vibrio for the open chromatin ATAC-Seq and histone ChIP-Seq (H3K4me3, H3K27ac, H3K27me3) samples.

**Supplementary Figure 2.** Genomic distribution of the ChIP-Seq high signal regions (pink) and low mappability regions of the turbot genome (blue) included in the turbot blacklist for each of the 22 chromosomes.

**Supplementary table 1.** Metadata of the **A)** 12 and 18 turbot specimens utilized for the in vitro (F) and in vivo (Fvv) challenges. **B)** 18 head kidney samples, extracted after immunostimulation, utilized for the *in vivo* challenges. **C**) 12 head kidney samples utilized for the *in vitro* challenges. **D**) 6 cell extracts pools (leukocytes), derived from the 12 head kidney samples utilized for the *in vitro* challenges. **E**) 18 primary cell cultures (leukocytes), sampled after immunostimulation and derived from the 6 leukocyte extracts utilized for the *in vitro* challenges.

**Supplementary table 2.** Metadata of the **A)** ENA accession IDs to the studies were the raw sequencing files used for the RNA-Seq, ATAC-Seq, ChIP-Seq and µChIPmentation analysis are allocated. **B)** ENA accession IDs, file names and md5 codes to the raw sequencing files used for the RNA-Seq, ATAC-Seq, ChIP-Seq and µChIPmentation analysis. **C)** General library metadata for the RNA-Seq, ATAC-Seq, ChIP-Seq and µChIPmentation analysis. Links to the detailed protocols, from sample homogenization to library size selection can be found in column M "Library Construction Protocol". **D)** Metadata for the RNA-Seq libraries. Links to the detailed extraction protocol, and library preparation can be found in columns K "Extraction Protocol" and AF "Library Generation Protocol", respectively. **E)** Metadata for the ChIP-Seq and µChIPmentation input libraries. Links to the detailed experimental protocol, and library preparation can be found in columns J "Experimental Protocol". **F)** Metadata for the ChIP-Seq and µChIPmentation libraries. Links to the detailed experimental protocol, and library preparation can be found in columns J "Experimental Protocol". **G)** Metadata for the ATAC-Seq libraries. Links to the detailed experimental protocol, and library preparation can be found in columns J "Experimental Protocol".

**Supplementary table 3.** Summary of the sequencing metrics of the **A)** RNA-Seq and **B)** ATAC-Seq and ChIP-Seq data, processed with the nf-core pipeline, of the *in vitro* and *in vivo* challenges.

**Supplementary table 4.** Differential gene expression results of the in vitro challenges stimulated with **A)** Poly (I:C) and **B)** Vibrio and in vivo challenge stimulated with **C)** Poly (I:C) and **D)** Vibrio (p-value < 0.05).

**Supplementary table 5.** GO enrichment analysis of the differentially expressed genes **A)** downregulated and **B)** upregulated for Poly (I:C) *in vitro*; **C)** downregulated and **D)** upregulated for *Vibrio in vitro*; **E)** downregulated and **F)** upregulated for Poly (I:C) *in vivo*; and **G)** downregulated and **H)** upregulated for Vibrio in vivo, for Biological Process (p-value < 0.05).

**Supplementary table 6. A)** Summary of the different GO analyses performed on the lists of common and exclusive differentially expressed genes (DEGs) for each comparison. GO enrichment on Biological Process was chosen as the default analysis (ShinyGO). If no significant enriched GO terms were identified, GO profiling on Biological Process was used (g:Profiler). If neither significant GO enrichment nor profile was found, GO terms for Biological Process for the DEGs in the list were obtained (ENSEMBL). **B)** Significantly enriched GO terms (Biological Process) for the different comparisons between DEGs. **C)** Functional profiling of significant GO terms (Biological Process) for the different comparisons between differentially expressed genes (DEGs) where no significant enrichment was found. **D)** GO terms (Biological Process) associated with the differentially expressed genes (DEGs) arisen from different comparisons between DEGs, where neither significant enrichment nor profile was found.

**Supplementary table 7.** ChIP-Seq and ChIPmentation turbot blacklist of low mappability and high input signal regions.

**Supplementary table 8. A)** Chromatin state annotation for the *in vitro* and *in vivo* challenges. Genome wide chromatin state distribution of the **B)** Control *in vitro*, **C)** Poly (I:C) *in vitro*, **D)** Vibrio *in vitro*, **E)** Control *in vivo*, **F)** Poly (I:C) *in vivo* and **G)** *Vibrio in vivo*.

**Supplementary table 9. A)** Differentially accessible regions (DAR) respective to the corresponding controls for the ATAC-Seq assays (p-value < 0.05). Differentially histone modification regions (DHMR) respective to the corresponding controls for the ChIP-Seq and µChIPmentation assays for the **B)** H3K4me3, **C)** H3K27ac and **D)** H3K27me3 marks (p-value < 0.05).

**Supplementary table 10.** Differential Accesibility Regions (DAR) and Differential Histone Modification Regions (DHMR) located in promoter regions and integrated with Differential Expressed Genes (DEG) for the in vivo and in vitro challenges stimulated with Poly (I:C) and Vibrio (Hypergeometric test, p-value < 0.05)

**Supplementary table 11.** GO enrichment analysis (Biological Process) for the upregulated differentially expressed genes (DEG) with differential accessibility regions (DAR) and/or differential histone modification regions (DHMR) in their promoters (p-value < 0.05).

**Supplementary table 12.** Transcription factor (TF) motif enrichment results for the differential accessibility regions (DAR) and differential histone modification regions (DHMR) in **A)** promoter regions and **B)** differential enhancer state-related regions in intergenic or intron regions. Only motifs with p-value < 0.01 and present in more than 5% of the target sequences were considered.

**Supplementary table 13.** Upregulated TFs with binding motif enrichment predicted on DARs and DHMRs (H3K4me3 and H3K27ac) of active promoters and enhancer regions. The second column shows TF with promoters overlapping with DAR/DHMR, while the third column shows TF that are differentially expressed genes (DEGs). TF that are in both columns are underlined.
